# HOXA10-TWIST2 Antagonism Drives Partial Epithelial-to-Mesenchymal transition for Embryo Implantation

**DOI:** 10.1101/2025.01.10.631632

**Authors:** Nancy Ashary, Sanjana Suresh, Anshul Bhide, Sharmishtha Shyamal, N Pranya, Anuradha Mishra, Saee Patil, A Anuradha, Shruti Hansda, B V Harshavardhan, Mohit Kumar Jolly, Deepak Modi

## Abstract

In mammalian reproduction, a significant proportion of embryos fail to implant despite a receptive uterus, suggesting that defects in epithelial remodelling at the embryo-uterine interface contribute to implantation failure. The molecular programs enabling such remodelling remain incompletely understood. Here, we identify a conserved transcriptional circuit involving HOXA10 and TWIST2 that regulates epithelial plasticity in the endometrium via partial epithelial-to-mesenchymal transition (pEMT). HOXA10, a transcription factor essential for uterine receptivity, is specifically downregulated in the luminal epithelium at implantation in mice, hamsters, and monkeys. Integrated CUT&RUN and transcriptomic profiling in human endometrial epithelial cells reveal that HOXA10 directly activates epithelial gene networks and represses mesenchymal programs. HOXA10 loss, both *in vitro* and *in vivo*, induces a pEMT state with increased cell motility. Mechanistically, HOXA10 represses TWIST2, a core EMT regulator; its derepression promotes mesenchymal gene expression and epithelial cell displacement. TWIST2 knockdown restores epithelial identity and impairs implantation. These findings establish a mutually antagonistic HOXA10-TWIST2 circuit as a key regulator of pEMT and epithelial remodelling during implantation.

**Graphical Abstract:** 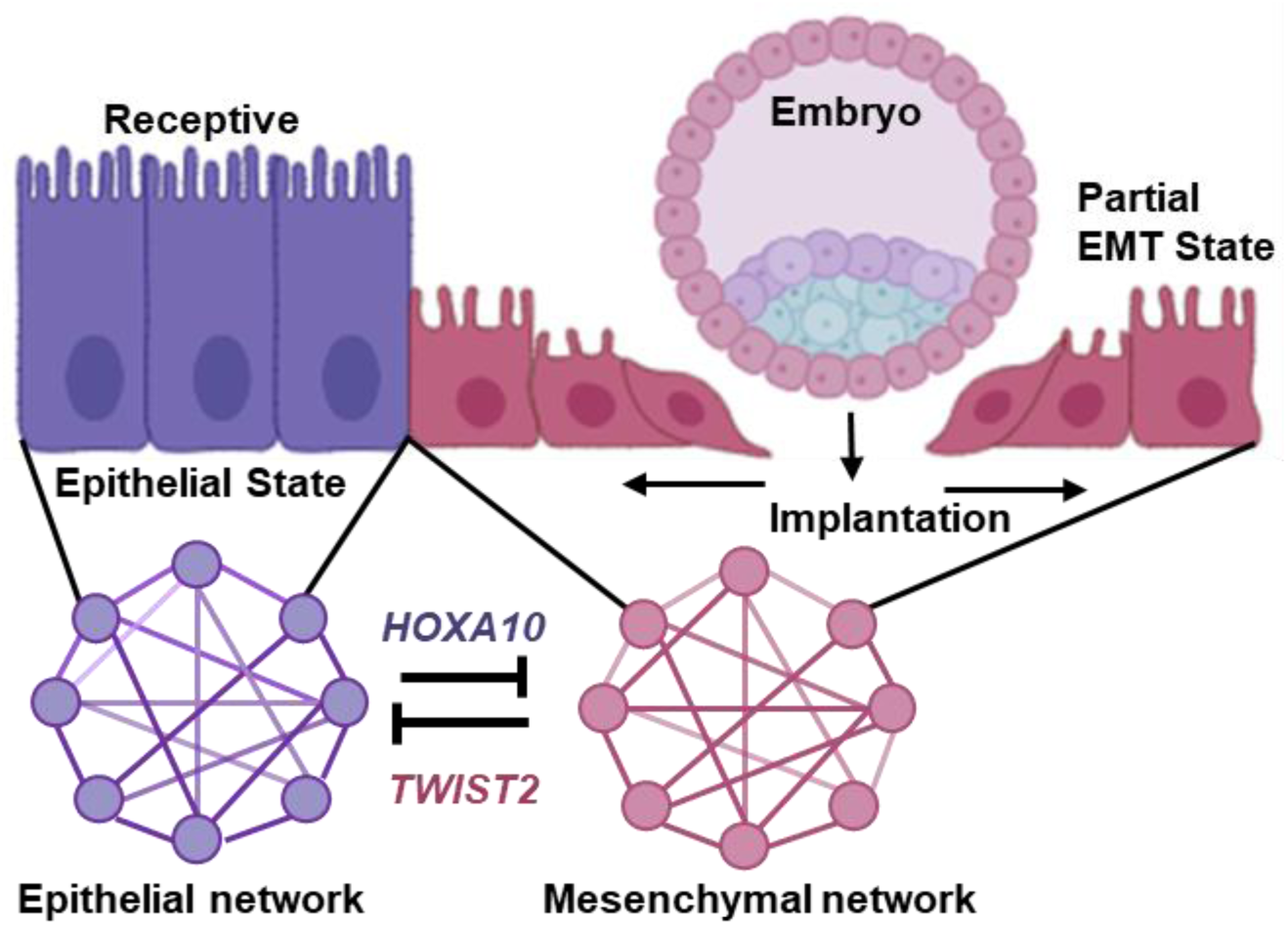

## Introduction

Tissue remodelling in adult organs requires a delicate balance between maintaining epithelial integrity and allowing transient plasticity. This balance is central to a variety of physiological processes, including wound healing and organ regeneration and also in pathological conditions like cancer progression and fibrosis (Pinet & McLaughlin, 2019; Talbott *et al*, 2022; Maggiore & Zhu, 2024). One of the most tightly regulated physiological examples of such remodelling is embryo implantation, where the endometrial epithelium transiently transforms to accommodate the embryo (Ashary *et al*, 2018; Ochoa-Bernal & Fazleabas, 2020). These processes mechanistically resemble those seen in cancer, fibrosis, and wound healing, making it a powerful model to understand regulated epithelial plasticity.

In mammals, embryo implantation marks the critical interface between maternal and embryonic tissues, establishing pregnancy (Ashary *et al*, 2018; Deng & Wang, 2022). Although the endometrium must acquire a receptive state to allow implantation, the failure of high-quality euploid embryos to implant in human assisted reproduction failure points to underlying and often unrecognized endometrial defects as significant contributors to early pregnancy loss (Dimitriadis *et al*, 2020). Thus, understanding the fundamental mechanisms that drive embryo implantation is crucial to improving the success of assisted reproduction. Implantation begins with the apposition and invasion into the underlying stroma (Ruane *et al*, 2022; Muter *et al*, 2023; Shibata *et al*, 2024). This process involves extensive remodelling of the epithelial barrier where the luminal epithelium undergoes structural and functional changes, including loss of apical-basal polarity, loosening of intercellular junctions, and reorganization of the cytoskeleton (Hochfeld *et al*, 1990; Thie *et al*, 1995; Murphy, 2000; Su *et al*, 2012; Jha *et al*, 2006; Ye, 2020; Whitby *et al*, 2020). Such changes are reminiscent of epithelial plasticity described as epithelial-to-mesenchymal transition (EMT), a process by which epithelial cells acquire mesenchymal traits such as motility and plasticity (Yang *et al*, 2020). Although EMT has been extensively studied in cancer and fibrosis, its role in physiological contexts like implantation is poorly understood.

Traditionally, epithelial-to-mesenchymal transition (EMT) has been viewed as a binary process in which epithelial cells undergo a reversible but complete transition to a mesenchymal state. However, emerging evidence challenges this model and suggest that EMT is not a linear trajectory but rather involves the acquisition of distinct, stable intermediate states that retain both epithelial and mesenchymal features. These states are collectively referred to as hybrid or partial EMT (pEMT) (Norgard *et al*, 2021; Subbalakshmi *et al*, 2022; Hari *et al*, 2022). pEMT is now recognized as a dominant mechanism in cancer progression and metastasis, where it is often dysregulated and persistent (Pastushenko & Blanpain, 2019). In contrast, the epithelial remodelling that occurs during embryo implantation may likely represent a physiological form of pEMT, which is tightly regulated, spatially confined, and reversible. However, whether pEMT occurs in the endometrium at the time of implantation remains unknown. Further, how such epithelial plasticity is transcriptionally orchestrated in adult tissues under physiological conditions is an unresolved but compelling question.

HOXA10, a highly conserved transcription factor of the Abd-B homeobox family, is a well-established regulator of uterine development, endometrial receptivity, and decidualization (Du & Taylor, 2016; Ashary *et al*, 2020; Bi *et al*, 2022). It is expressed in both the epithelial and stromal compartments of the endometrium and is indispensable for implantation (Benson *et al*, 1996). Also, aberrant HOXA10 expression is associated with several uterine pathologies, including endometriosis, endometrial cancer, and recurrent implantation failure (Zanatta *et al*, 2010; Mishra *et al*, 2022; Mishra & Modi, 2023, 2024; Pîrlog *et al*, 2025). Despite its indispensable role in implantation, we and others have observed that while HOXA10 expression peaks during the receptive phase, it is selectively downregulated in the luminal epithelium at the site of embryo implantation (Godbole *et al*, 2007; Guo *et al*, 2009). This observation appears paradoxical, considering its essential role in establishing uterine receptivity. However, considering its dynamic regulation both spatially and temporally, we hypothesized that the downregulation of HOXA10 may be functionally significant and could serve as a molecular switch that permits embryo implantation.

Here, we demonstrate that HOXA10 maintains epithelial identity in the endometrial epithelium by activating epithelial gene programs and repressing mesenchymal ones. Its local loss at implantation sites triggers pEMT and cell migration, facilitating epithelial remodelling and embryo embedding. Mechanistically, we identify TWIST2, a known EMT transcription factor, as a direct target of HOXA10. TWIST2 derepression promotes mesenchymal gene expression and cellular motility, while its knockdown restores epithelial features and impairs implantation. These findings establish embryo implantation as a physiological model of pEMT, driven by a precise transcriptional circuit, and reveal how adult tissues transiently alter identity to enable specialized functions.

## Results

### HOXA10 is Downregulated in the Endometrial Luminal Epithelium at the Time of Embryo Implantation

Previous studies have shown that HOXA10 is downregulated in the luminal epithelium of the endometrium at the time of implantation in monkeys and canines (Godbole *et al*, 2007; Guo *et al*, 2009). To determine whether a similar downregulation occurs in rodents, we assessed HOXA10 expression in the endometrium of mice and hamsters. In mice, HOXA10 was abundantly expressed in both the cytoplasm and nuclei of luminal epithelial cells during diestrus and on day 4 (D4) of pregnancy, when the endometrium is receptive (Fig. 1A). On the morning of day 5 (D5), following embryo attachment, HOXA10 in the luminal epithelial cells decreased. By the evening of D5, HOXA10 in most epithelial cells was reduced. Compared to D4, HOXA10 levels declined by ∼60% on the morning of D5 (09:00h), and by ∼80% in the evening (21:00h); the reduction was statistically significant at both time points (Fig. 1B). This downregulation was restricted to implantation sites, as luminal epithelial cells at inter-implantation sites on D5 retained HOXA10 expression at levels similar to those on D4 (Fig. S1).

**Fig. 1:**
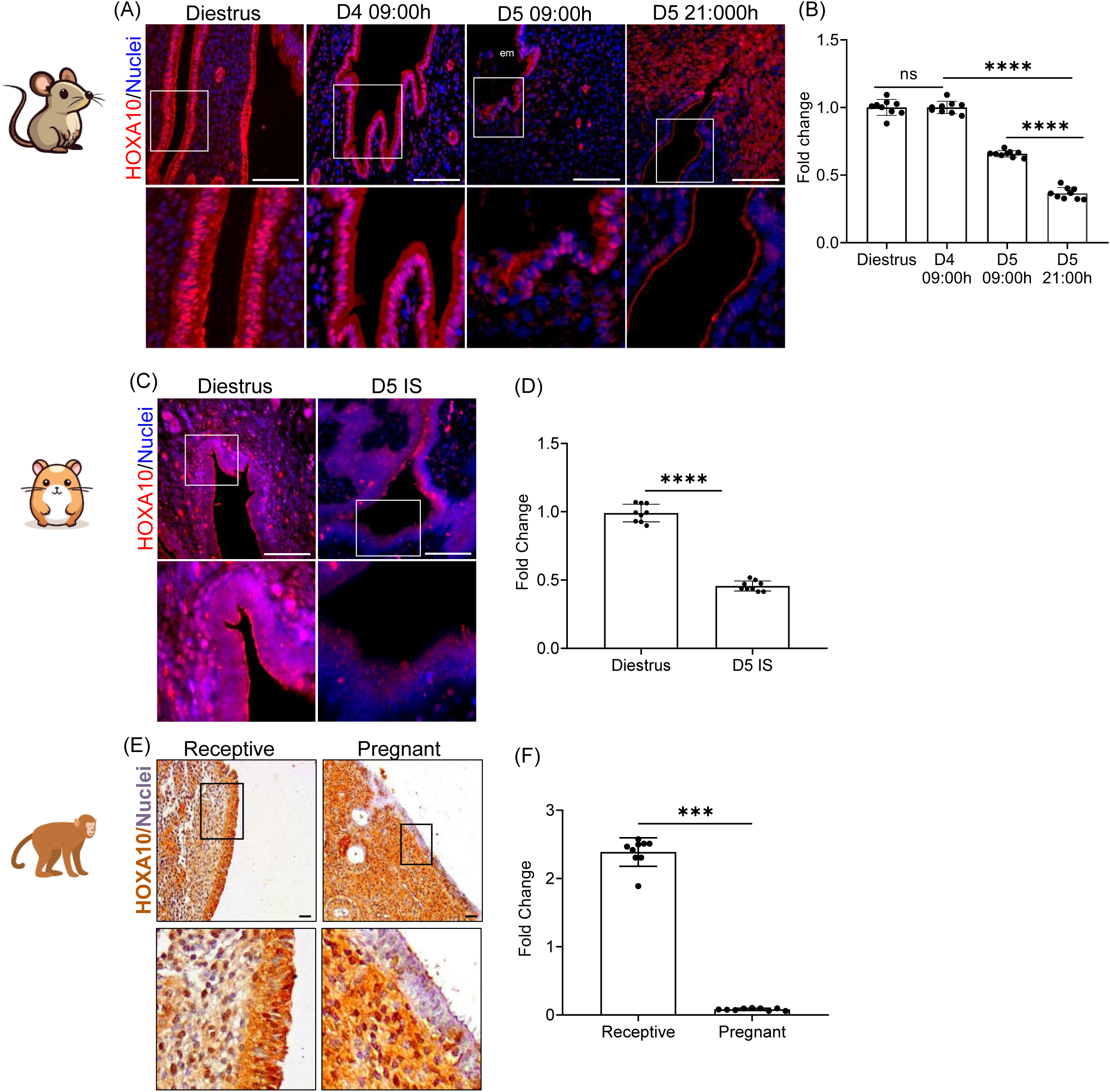
HOXA10 is Downregulated in the Endometrial Luminal Epithelium at the Time of Embryo Implantation. **(A)** Immunofluorescence for HOXA10 in the mouse uterus of diestrus stage, D4 09:00h, D5 09:00h and D5 21:00h. Selected area is boxed and zoomed. **(B)** Quantification of immunostaining for HOXA10 in the mouse uterus. Each dot represents fluorescence intensity in a single region of interest and data of 3 independent regions from multiple sections of 3 animals are plotted (n=3). **(C)** Immunofluorescence for HOXA10 in hamster uterus of diestrus stage, D5 Implantation site (IS). Selected area is boxed and zoomed. **(D)** Quantification of HOXA10 signals in the hamster uterus. Each dot represents fluorescence intensity in a single region of interest and data of 3 independent regions from multiple sections of 3 animals are plotted (n=3). **(E)** Immunohistochemistry for HOXA10 in the bonnet monkey uterus of receptive stage and pregnant. **(F)** Quantification of immunostaining for HOXA10 in the bonnet monkey uterus. Each dot represents fluorescence intensity in a single region of interest and data of 3 independent regions from multiple sections of 3 animals are plotted (n=3). em = embryo, Scale bar = 100µm, ns is non-significant, **** is p<0.0001 and *** is p<0.001 with mean ±SD values are shown

In hamster endometrium, HOXA10 was also abundantly expressed in luminal epithelial cells during diestrus. Compared to non-pregnant controls (diestrus), HOXA10 expression was significantly reduced at implantation sites on D5 by nearly 50% (Fig. 1C and D). Similar to mice, this reduction was restricted to implantation sites, as the inter-implantation sites had HOXA10 expression similar to non-pregnant controls (Fig. S2).

We revisited our HOXA10 immunostaining data in the bonnet monkey endometrium and quantified the staining intensities. In the receptive phase, HOXA10 was strongly expressed in both the cytoplasm and nuclei of luminal epithelial cells. However, in animals that were mated and the presence of the embryo verified (Rosario *et al*, 2005), HOXA10 expression was markedly diminished in luminal epithelial cells (Fig. 1E). Quantification revealed an ∼80% reduction in HOXA10 levels in these animals compared to the receptive phase (Fig. 1F).

These findings together indicate that embryo implantation is associated with site-specific downregulation of HOXA10 in the endometrial luminal epithelium, and this phenomenon is conserved across rodents and primates.

### Loss of HOXA10 Induces Migration of Endometrial Epithelial Cells

To understand the significance of the loss of HOXA10 in the endometrial epithelial cells, we stably knocked down its expression in the human endometrial epithelial cell line and performed RNAseq (Fig. 2A). As compared to the scrambled shRNA expressing cells (control), in the cells expressing *HOXA10* shRNA (*HOXA10*KD), there was a 70% reduction in *HOXA10* mRNA (Fig. S3). Transcriptome analysis of *HOXA10*KD cells revealed significant alterations in gene expression, where 2148 genes were upregulated and 2915 genes were downregulated as compared to control cells, highlighting the substantial impact of *HOXA10* knockdown on endometrial gene regulation (Fig. 2B and C). We performed Gene Set Enrichment Analysis (GSEA) of the differentially expressed genes (DEGs) and observed a significant enrichment of biological processes associated with tissue migration, actin filament-based movement, cell-substrate adhesion, and cell junction assembly (Fig. 2D) The molecular functions, cellular components and pathways associated with the DEGs also revealed a significant enrichment of processes associated with cell adhesion, cytoskeletal organization, and motor proteins (Fig. S4). These observations prompted us to hypothesize that loss of HOXA10 may alter the migration of endometrial epithelial cells. To address this, we performed live cell imaging of control and *HOXA10*KD endometrial epithelial cells and observed that the control cells were largely stationary while most of the *HOXA10*KD cells were migratory (Fig. 2E, Movie S1 and S2). In contrast to control cells that were hardly displaced, the *HOXA10*KD cells showed large displacement and migrated with a higher velocity than controls (Fig. 2F and G). The increase in the migratory property of the *HOXA10*KD cells was also associated with altered actin filament organization. In the control cells, the F-actin staining was largely cytoplasmic with a few organized in cortical bundles. In contrast, in the *HOXA10*KD cells, F-actin assembled into thick, parallel bundles that ran perpendicular to the cell edge, and in most cells, a high number of F-actin-rich projections were observed (Fig. 2H). Both of these features are characteristic of migratory cells. These results indicate that loss of HOXA10 induces migration of endometrial epithelial cells.

**Fig. 2:**
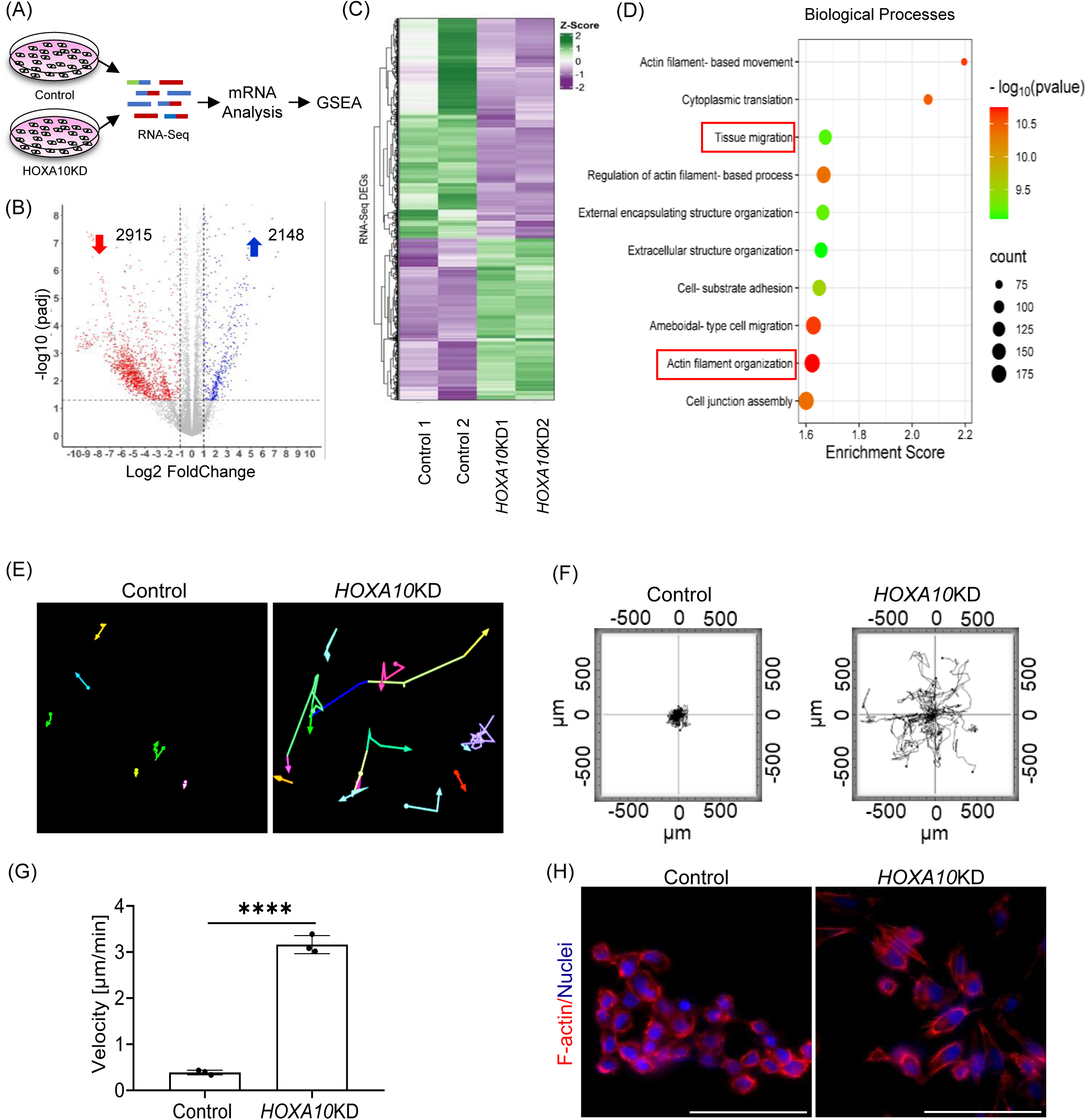
Loss of HOXA10 Induces Migration of Endometrial Epithelial Cells. **(A)** Schematic representation of RNA-seq experimental set up. **(B)** Volcano plot showing the differentially expressed genes between *HOXA10*KD and control. **(C)** Heatmap depicts the Z-score normalized expression levels of RNA-Seq DEGs in two control samples (Control 1 and Control 2) and two HOXA10 knockdown samples (*HOXA10*KD1 and *HOXA10*KD2) cells. Green and purple colours represent upregulated and downregulated genes, respectively, relative to the average expression across all samples. Hierarchical clustering was performed to identify distinct expression patterns. **(D)** Top enriched biological processes associated with the differentially expressed genes (DEGs). **(E)** Stable lines of human endometrial epithelial cells (RL95-2) expressing scrambled shRNA (Control) and HOXA10 shRNA (*HOXA10*KD) were subjected to live imaging. Tracks of control and *HOXA10*KD cells for 24h (movie S1 and S2). **(F)** Representative rose plots depicting the trajectories of control (n=22 cells) and *HOXA10*KD (n=22 cells) endometrial epithelial cells are also shown. **(G)** Quantification of migration velocity of control and *HOXA10*KD endometrial epithelial cells (n=3). **(H)** Immunostaining for F-actin in control and *HOXA10*KD endometrial epithelial cells (n=3). Scale bar = 100µm. **** is p<0.0001 with mean ±SD values are shown.

### Loss of HOXA10 Aids in Epithelial Cell Displacement for Embryo Implantation

Given that loss of HOXA10 induced epithelial cell migration, we asked whether it facilitates their displacement during embryo implantation. To test this, we modified the *in vitro* embryo implantation assay (Thie *et al*, 1997),where instead of counting the number of attached spheroids, we assessed the extent of displacement of epithelial cells surrounding the spheroids. Fluorescently labelled trophoblast spheroids were cultured on confluent monolayers of fluorescently labelled control and *HOXA10*KD cells. After 24 hours, the displacement of epithelial cells surrounding the spheroids was tested (Fig. 3A). In the controls, minimal clearance of the epithelial monolayer was observed beneath the trophoblast spheroid. In contrast, *HOXA10*KD exhibited a pronounced zone of clearance, with the region beneath and around the spheroid largely devoid of epithelial cells (Fig. 3B). As there were no trophoblast cells outside the spheroid, we concluded that this clearing is due to the migration of the epithelial cells and not a displacement by the trophoblast cells. Indeed, live cell imaging revealed that within 6 hours of co-culture, *HOXA10*KD epithelial cells adjacent to the spheroid became motile and migrated away, resulting in a substantial zone of clearance by 12 hours (Fig. 3C, Movie S3).

**Fig. 3:**
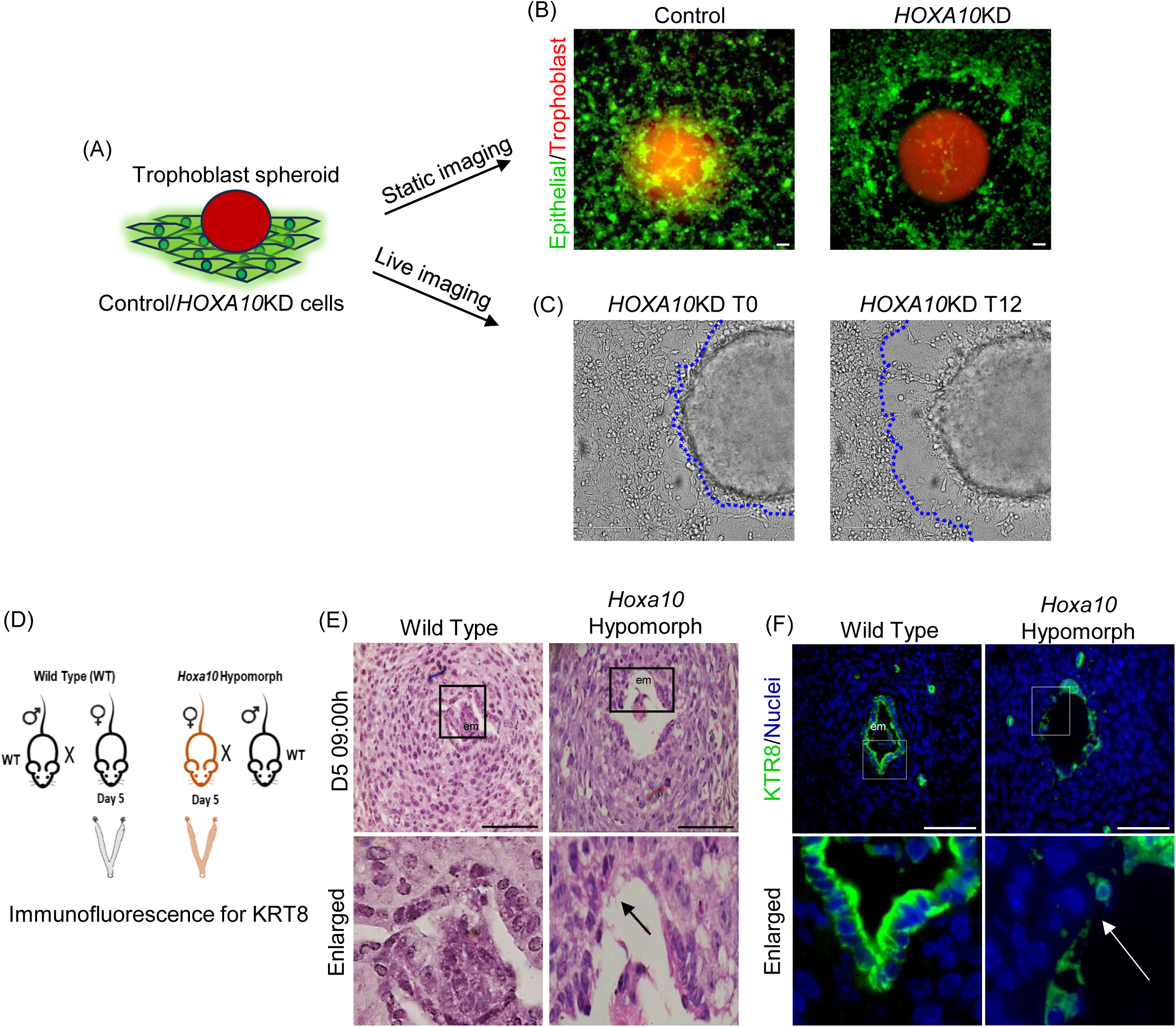
Loss of HOXA10 Aids in Epithelial Cell Displacement for Embryo Implantation. **(A)** Schematic representation of experimental set up. Trophoblast spheroids were labelled with CellTracker Red CMTPX and co-cultured with monolayers of CellTracker Green CMFDA labelled control or *HOXA10*KD cells (n=3). **(B)** Displacement of endometrial epithelial cells in response to trophoblast spheroid. Monolayers of control and *HOXA10*KD cells were incubated with trophoblast spheroid and imaged after 24h. **(C)** Time-lapse of *HOXA10*KD cells incubated with trophoblast spheroid (movie S3). Static images at T0 (Time Zero) and T12h are shown. Marked are the boundaries of the epithelial cell layer. **(D)** Schematic representation of experimental set up. Mouse uteri of wildtype and *Hoxa10* hypomorph on D5 09:00h were collected. **(E)** Haematoxylin and Eosin-stained sections of wildtype and *Hoxa10* hypomorphs mouse uterus on D5 09:00h (n=3). **(F)** Immunofluorescence for KRT8 in wildtype and *Hoxa10* hypomorphs uteri on D5 09:00h (n=3). The selected area is boxed and zoomed. em = embryo. Scale bar = 100µm.

To determine if HOXA10 loss also promotes epithelial cell displacement i*n vivo* during implantation, we examined implantation crypts from *Hoxa10* hypomorphs. *Hoxa10* hypomorphs are transgenic mice expressing a shRNA targeting *Hoxa10*, and have significantly reduced HOXA10 levels in the endometrium (Mishra *et al*, 2022), including the luminal epithelium (Fig. S5). We hypothesized that if loss of HOXA10 induces migration, the luminal epithelial layer in hypomorphs would exhibit premature displacement within the implantation crypt. Previous studies have shown that on the morning of day 5 of mouse pregnancy, the implantation crypt is formed, and the embryo is enclosed by an intact luminal epithelial layer. By the evening of D5, breaching is initiated, and luminal epithelial cells begin to displace (Li *et al*, 2015). Therefore, we analysed the epithelial lining in the implantation crypt of wild-type and *Hoxa10* hypomorphs on D5 morning (Fig. 3D). As expected, in the wild-type animals the implantation sites had an intact luminal epithelium with a continuous layer of KRT8-positive cells (Fig. 3E and F). In contrast, *Hoxa10* hypomorphs exhibited gaps in the luminal epithelium similar to those seen in wild-type D5 evening samples and KRT8-positive cells appeared displaced (Fig. 3E and F). These findings support that HOXA10 loss promotes luminal epithelial cell motility, facilitating their displacement during embryo implantation.

### HOXA10 Occupies Genomic Sites Co-regulated by Implantation and Epithelial Plasticity Transcription Factors

The observation that HOXA10 knockdown promotes epithelial cell migration and displacement (Fig. 2E and 3B) suggested that it might play a broader role in maintaining epithelial identity. To explore this possibility, we mapped the genome-wide chromatin occupancy of HOXA10 in endometrial epithelial cells using CUT&RUN with a validated antibody (Fig. S6) and identified 69,990 high-confidence binding sites. Analysis of the top 10,000 peaks revealed that most HOXA10 binding events occurred in intergenic and intronic regions, with a subset mapping to promoters and transcription start sites (TSSs) (Fig. S7). Aggregate peak plots confirmed strong enrichment at these sites (Fig. 4A).

**Fig. 4:**
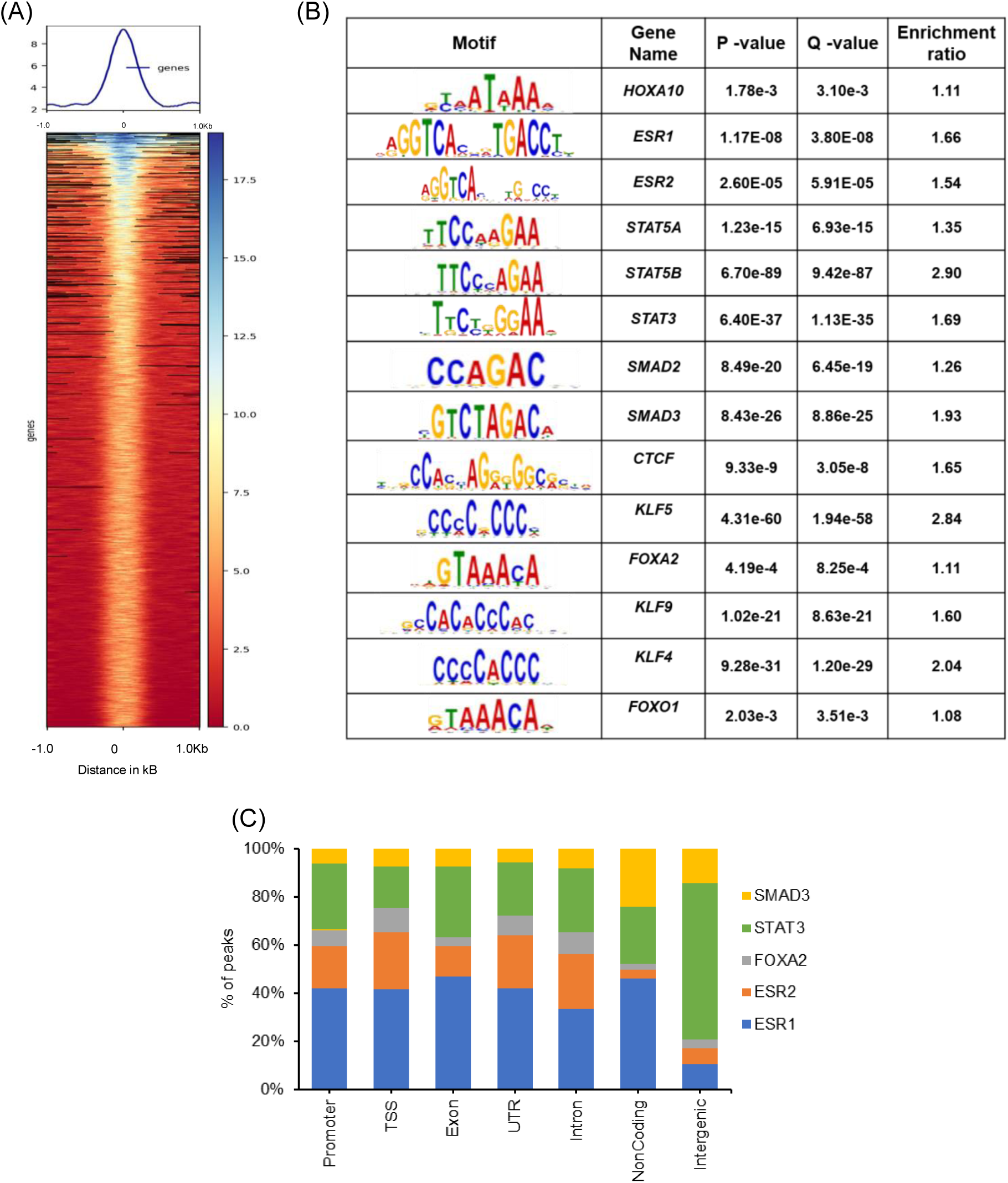
HOXA10 Occupies Genomic Sites Co-regulated by Implantation and Epithelial Plasticity Transcription Factors. **(A)** The heat map and profile plot display the region occupied by HOXA10 in the endometrial epithelial cells. **(B)** Non-HOX motifs enriched in HOXA10 CUR&RUN. **(C)** Percentage of HOXA10 co-enriched gene motif overlapping peak

Motif analysis of the top 5,000 peaks revealed a significant enrichment for binding motifs of multiple transcription factors, including CTCF, a finding consistent with previous datasets from chicken cells, (Jerković *et al*, 2017) hence validating our approach. Amongst the significantly co-enriched motifs, many belonged to transcription factors known to regulate uterine receptivity and embryo implantation, such as ERα (ESR1), ERβ (ESR2), STAT3, STAT5A/B, SMAD2, SMAD3, FOXA2, FOXO1, KLF4, KLF5, and KLF9 (Fig. 4B;Table S1). Notably, HOX motifs are also enriched in ERα ChIP-seq datasets from estrogen-treated mouse uterus, suggesting that HOXA10 and estrogen receptors may co-regulate gene expression programs relevant to implantation (Hewitt *et al*, 2012). In our dataset, ERα and ERβ motifs were predominantly localized to promoters, TSSs, untranslated regions (UTRs), and exons, indicating their shared regulatory role with HOXA10 (Fig. 4C). Given the essential role of estrogen signalling in the uterine epithelium (Pawar *et al*, 2015; Robertshaw *et al*, 2016). Our results support the idea that HOXA10 acts as part of a larger transcriptional network governing epithelial function.

To understand the biological significance of this network, we examined HOXA10-bound genes enriched at promoter and TSS regions. Pathway analysis revealed associations with signalling cascades central to implantation, including VEGF, AMPK, Hippo, and TGF-β pathways (Fig. S7), all of which have previously been linked to embryo implantation (Table S2) and epithelial plasticity (Nieto & Cano, 2012; Yan *et al*, 2024).

Together, these findings suggest that HOXA10 regulates a diverse set of genes through its direct occupancy of regulatory elements, functioning in coordination with multiple transcription factors involved in implantation and epithelial remodelling. The enrichment of motifs for regulators of epithelial plasticity, including SMADs, FOXOs, and KLFs led us to next investigate whether HOXA10 may directly influence EMT programs in the endometrial epithelium.

### HOXA10 Directly Regulates Genes Involved in Epithelial-to-Mesenchymal Transition

To identify direct transcriptional targets of HOXA10, we intersected the DEGs from *HOXA10*KD cells with genes bearing HOXA10 occupancy at promoter or TSS regions. Of the 5,378 high-confidence HOXA10 peaks located at gene regulatory sites, 1,246 genes (24.6%) were differentially expressed upon loss of HOXA10 (Fig. 5A). These genes were designated as direct targets of HOXA10.

**Fig. 5:**
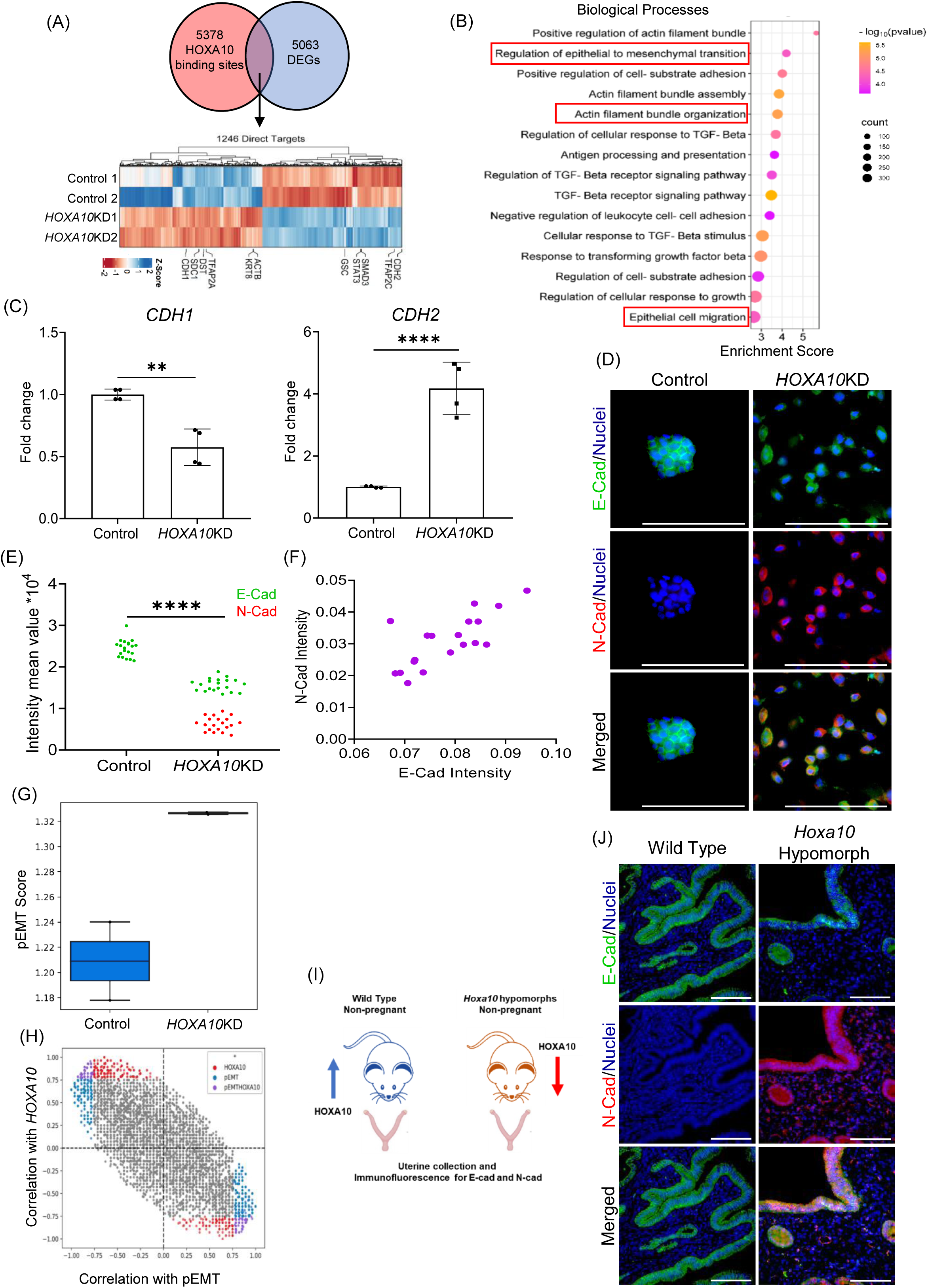
HOXA10 Direct Targets have a Role in Epithelial-to-Mesenchymal Transition (EMT). **(A)** Direct targets of HOXA10 were identified as genes with promoters/TSS with HOXA10 occupancy and having differential expression in HOXA10 knockdown (*HOXA10*KD) RL95 cells. Heatmap depicts the expression of the direct target of HOXA10 in the control and *HOXA10*KD cells. **(B)** Top enriched biological processes associated with the differentially expressed genes (DEGs). **(C)** mRNA levels of *CDH1* and *CDH2* (normalize to 18s) in RL95 cells stably expressing scrambled shRNA (Control) and *HOXA10* shRNA (*HOXA10KD)*. Y-axis is a fold change where values obtained from control cells were taken as 1. Data is the mean ± SD for the four independent replicates. **(D)** Co-immunostaining for E-Cad and N-Cad in control and *HOXA10*KD endometrial epithelial cells (RL95). **(E)** Quantification for E-Cad and N-Cad in control and *HOXA10*KD endometrial epithelial cells. Each dot represents fluorescence intensity in a single region of interest and data of 4 independent replicates are plotted. **(F)** Quantification of co-expression of E-Cad and N-Cad in single cell of HOXA*10*KD. Each dot represents fluorescence intensity in a single cell and data of 20 cells are plotted. **(G)** pEMT scores (mean ±SD) computed from the transcriptomes of *HOXA10*KD and control cells. **(H)** Scatter plot of Pearson correlations between a gene and pEMT score versus its correlation with HOXA10 levels. Genes correlated with HOXA10, pEMT, or both are highlighted in red, blue, and purple, respectively. **(I)** Schematic representation of experimental set up. Mouse uteri of non-pregnant wildtype and *Hoxa10* hypomorph were collected. **(J)** Co-immunostaining for E-Cad and N-Cad in wildtype and *Hoxa10* hypomorph mouse uterus at diestrus stage. Scale bar = 100µm. **** is p<0.0001 and ** is p<0.005 with mean ±SD values are shown.

Expression profiling of these direct targets revealed both upregulated and downregulated subsets (Fig. 5A), suggesting that HOXA10 functions as both an activator and repressor of transcription. GSEA of the upregulated direct targets revealed significant enrichment of pathways linked to EMT, actin remodelling, epithelial cell migration, and TGF-β signalling (Fig. 5B), with similar enrichment trends seen at the level of cellular processes and pathways (Fig. S8).

Given that *HOXA10* knockdown induced cell migration and that its direct targets are enriched for EMT-related functions, we next asked whether loss of HOXA10 promotes an EMT-like phenotype. Indeed, qPCR revealed a significant downregulation of *CDH1* (E-Cadherin) and a marked upregulation of *CDH2* (N-Cadherin) in *HOXA10*KD cells as compared to controls (Fig. 5C). Immunofluorescence confirmed that control epithelial cells expressed high levels of E-Cadherin and lacked N-Cadherin, whereas *HOXA10*KD cells showed reduced E-Cadherin and increased N-Cadherin expression (Fig. 5D and E). Notably, all *HOXA10*KD cells co-expressed E-Cadherin and N-Cadherin at varying levels (Fig. 5F) an expression pattern consistent with pEMT.

pEMT is a hybrid state where cells retain epithelial features, such as cell-cell adhesion, while also acquiring mesenchymal traits, including increased motility. This state is also marked by co-expression of epithelial and mesenchymal markers and is distinct from full EMT. We and others have demonstrated that such partial EMT states could be captured using transcriptome-derived scores (Puram *et al*, 2017; Chakraborty *et al*, 2020). Applying this approach to RNA-seq data from *HOXA10*KD cells, and in agreement with our immunostaining results, we observed significantly elevated pEMT scores compared to controls (Fig. 5G), indicating that loss of HOXA10 drives the cellular transcriptome to a pEMT state. To further probe the transcriptional relationship between HOXA10 and EMT, we computed Pearson correlation coefficients for all DEGs against both HOXA10 expression and pEMT scores. Genes that were positively correlated with HOXA10 were inversely correlated with pEMT scores, and vice versa (Fig. 5H), suggesting that HOXA10 acts as a stabilizer of the epithelial state. Its loss appears to derepress mesenchymal programs and shift cells toward pEMT.

To determine whether this transition also occurs *in vivo*, we analysed the uterine luminal epithelium of non-pregnant *Hoxa10* hypomorphic mice (Fig. 5I). In wild-type animals, the luminal epithelial cells had robust E-Cadherin expression with no detectable N-Cadherin. In contrast, luminal epithelial cells of *Hoxa10* hypomorphs co-expressed E-Cadherin and N-Cadherin, consistent with a pEMT state (Fig. 5J; Pearson correlation = 0.7). These findings suggest that HOXA10 is necessary to maintain epithelial identity in the uterus and that its reduction is sufficient to trigger pEMT *in vivo*.

### The Luminal Epithelium Undergoes pEMT at the Time of Embryo Implantation

Since HOXA10 expression naturally decreases in the luminal epithelium at implantation sites, and its loss promotes pEMT, we next investigated whether luminal epithelial cells undergo pEMT during the window of embryo implantation. We performed double immunofluorescence for E-Cadherin and N-Cadherin on uterine sections on D4 to D5 (midnight) in pregnant mice. On D4 09:00h, when HOXA10 is highly expressed, luminal epithelial cells showed strong lateral E-Cadherin expression and no N-Cadherin signal (Fig. 6A). By D5 09:00h, when HOXA10 levels decline at the implantation site, E-Cadherin expression became more diffuse, and N-Cadherin was detectable in several cells. By D5 21:00h and D5 24:00h, nearly all luminal epithelial cells within the implantation crypt co-expressed both markers (Fig. 6A), with a Pearson correlation of 0.79.

**Fig. 6:**
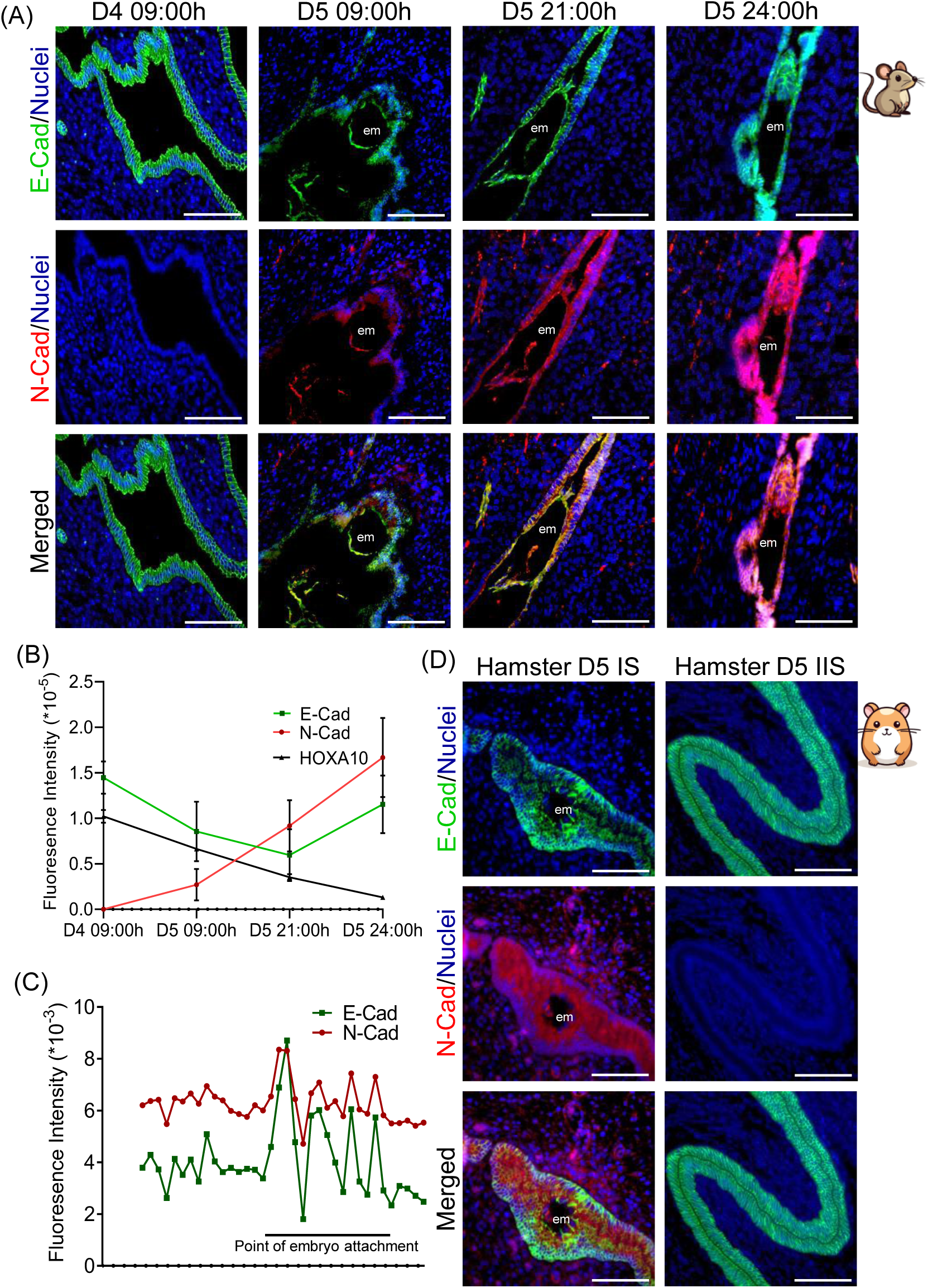
The Luminal Epithelial Cells are in pEMT State at the Time of Embryo Implantation. **(A)** Co-immunostaining for E-Cad and N-Cad in mouse uterus on D4 09:00h, D5 09:00h, D5 21:00h and D5 24:00h. **(B)** Quantification of immunostaining for HOXA10, E-Cad and N-Cad. values are mean ±SD and n=5 animals per group. **(C)** Spatial distribution of endometrial epithelial cells in different pEMT states at the site of embryo attachment on D5 21:00h. **(D)** Co-immunostaining for E-Cad and N-Cad in the hamster uterus on D5 at the implantation site (IS) and inter-implantation site (IIS). em = embryo, Scale bar = 100µm.

Quantitative analysis across timepoints confirmed a progressive decline in HOXA10 and E-Cadherin expression from D4 to D5, accompanied by a corresponding increase in N-Cadherin levels (Fig. 6B). Single-cell quantification at D5 21:00h showed that epithelial cells adjacent to the embryo consistently co-expressed E-Cadherin and N-Cadherin, while cells further from the embryo exhibited more variable levels of both markers (Fig. 6C).

To assess whether this phenomenon is conserved, we examined luminal epithelial cells in D5 pregnant hamsters. In inter-implantation regions, cells exclusively expressed E-Cadherin. However, at the implantation crypts, many cells co-expressed E-Cadherin and N-Cadherin (Fig. 6D; Pearson correlation = 0.5), indicating that pEMT is not only a feature of mouse implantation but also occurs in the hamster uterus.

Together, these results show that pEMT is a physiological feature of the luminal epithelium during embryo implantation, and that HOXA10 downregulation may act as a trigger for this transition.

### HOXA10 Maintains Epithelial Identity by Activating Epithelial Genes and Repressing Mesenchymal Programs

Building on our observations that HOXA10 loss induces a pEMT phenotype and increases epithelial cell motility, we next sought to dissect the underlying transcriptional programs driving this shift. We curated a list of 1,918 genes implicated in EMT from the dbEMT and EMTome databases (Zhao *et al*, 2019; Vasaikar *et al*, 2020). Strikingly, 678 (35%) of these were differentially expressed in *HOXA10*KD cells, indicating that HOXA10 exerts widespread regulatory control over EMT-associated genes (Fig. 7A). GSEA of these 678 HOXA10-regulated EMT genes revealed significant enrichment of biological processes such as tissue migration, mesenchyme development, cell-substrate adhesion, epithelial morphogenesis, and proliferation (Fig. 7B). These findings suggested that HOXA10 may orchestrate the balance between epithelial and mesenchymal gene programs to maintain epithelial homeostasis.

**Fig. 7:**
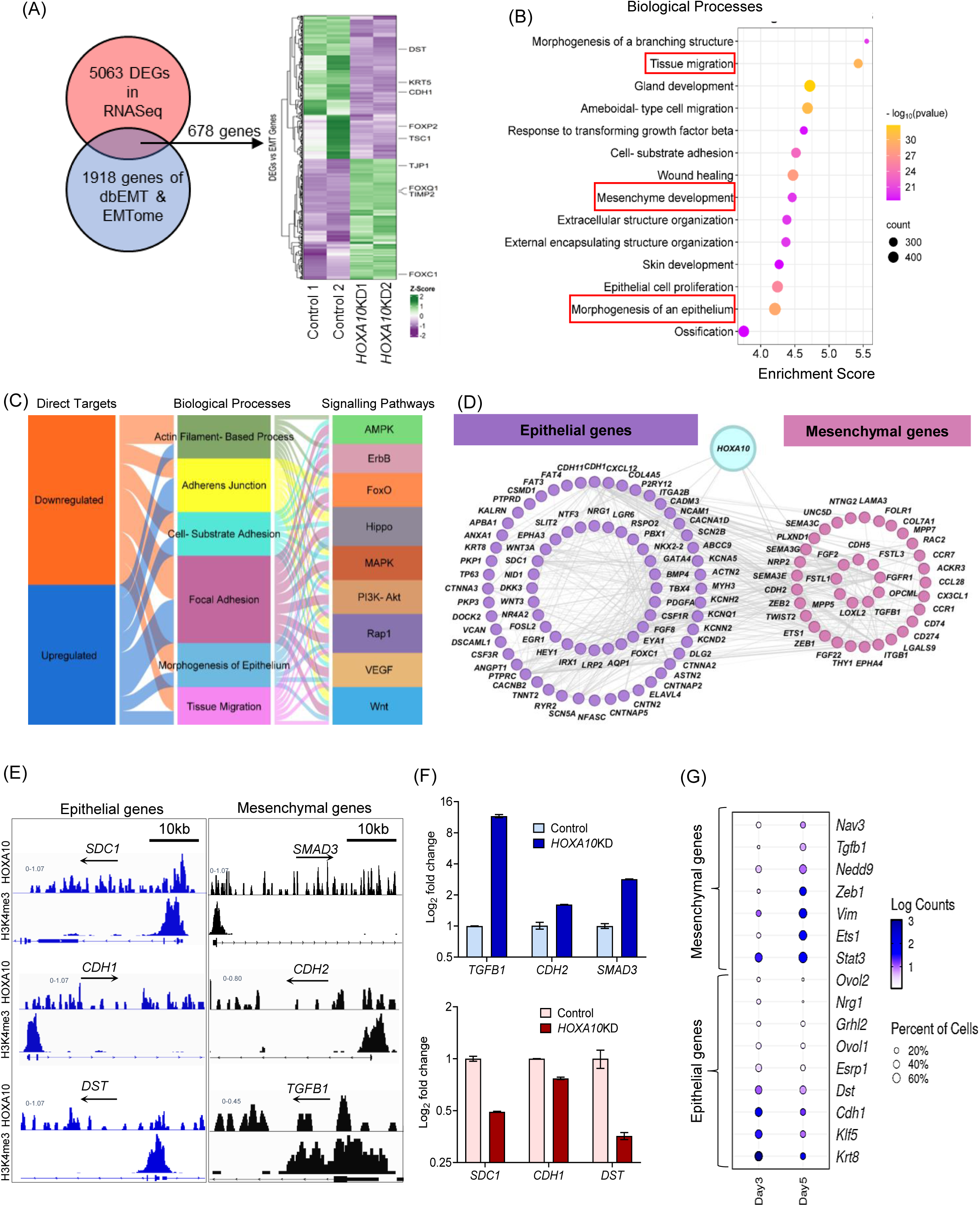
HOXA10 Maintains Epithelial Identity by Activating Epithelial Genes and Repressing Mesenchymal Programs. **(A)** Venn diagram for the DEGs and genes in dbEMT & EMTome datasets to identify the EMT-associated genes. Heatmap of 678 differentially expressed EMT-associated genes. **(B)** The biological processes significantly enriched with 678 differentially expressed EMT-associated genes. **(C)** Distribution of direct target of HOXA10 across enriched biological processes and pathways associated with EMT. **(D)** Protein-protein interaction (PPI) networks of the direct target of HOXA10 involved in epithelial and mesenchymal function. **(E)** Occupancy of HOXA10 in three epithelial genes (*SDC1, DST, CDH1*) and three mesenchymal genes (*TGFB1, CDH2, SMAD3*) are shown in IGV plots, H3K4Me3 marks the promoters of the same genes. The arrow depicts the direction of gene transcription. **(F)** mRNA levels of three epithelial and three mesenchymal genes (shown in panel E) in the control vs *HOXA10*KD cells. Values are mean ±SD of the fold change compared to controls. **(G)** Dot plot depicting gene expression of mesenchymal and epithelial genes in the luminal epithelium cells of mouse uteri on day 3 and day 5 of pregnancy. Data was extracted from single-cell RNA-seq of mouse uteri (Wang et al., 2023) [GSA: CRA011267].

To test this, we mapped the upregulated and downregulated HOXA10 direct target genes across hallmark EMT-related processes and KEGG pathways and observed that both groups of genes were evenly distributed across these categories, suggesting a broad impact of HOXA10 on both epithelial and mesenchymal gene networks (Fig. 7C). Indeed, Protein–Protein Interaction (PPI) network analysis revealed two prominent gene clusters: one composed of 24 genes involved in epithelial development and cell junction assembly, and another of 20 genes enriched for TGF-β signalling, cell motility, and canonical EMT regulators (Fig. 7D). Together with our transcriptomic correlation data (Fig. 5H), these findings point to a central role for HOXA10 in activating epithelial identity while repressing mesenchymal traits.

We next examined a subset of six HOXA10 direct target genes, three epithelial genes (*CDH1, SDC1, DST*) chosen based on their known expression in the human endometrial epithelium (Fig. S9) and three mesenchymal genes (*TGFB1, SMAD3, CDH2*) and strong CUT&RUN signals. IGV analysis confirmed HOXA10 occupancy at promoter and gene body regions for each of these targets (Fig. 7E). In *HOXA10*KD cells, the epithelial genes were significantly downregulated, while the mesenchymal genes were upregulated (Fig. 7F), supporting a model in which HOXA10 simultaneously sustains the epithelial program and prevents EMT.

To test whether this regulatory model holds true *in vivo*, we analysed published single-cell RNA-seq data from *Tacstd2*+ luminal epithelial cells on D3 and D5 of mouse pregnancy (Wang *et al*, 2023a). Compared to D3, epithelial HOXA10 targets were expressed at lower levels on D5, while mesenchymal targets showed increased expression (Fig. 7G). These shifts in gene expression, coinciding with natural HOXA10 downregulation at implantation sites, suggest that HOXA10 directly modulates epithelial plasticity during embryo implantation.

### Mutual Antagonism Between HOXA10 and TWIST2 Controls pEMT and Initiates Implantation

EMT is canonically regulated by transcription factors of the SNAI, ZEB, and TWIST families (Debnath *et al*, 2022). We interrogated the *HOXA10*KD gene regulatory network and identified key EMT regulators including *ZEB1, ZEB2,* and *TWIST2* within the differentially expressed gene set (Fig. 8A). Among these, *TWIST2* stood out not only as a DEG but also as a direct HOXA10 target, with its promoter and TSS region enriched for HOXA10 motifs (Fig. 8B and C). This suggested a potential regulatory axis in which HOXA10 directly represses TWIST2, a known driver of mesenchymal programs.

**Fig. 8:**
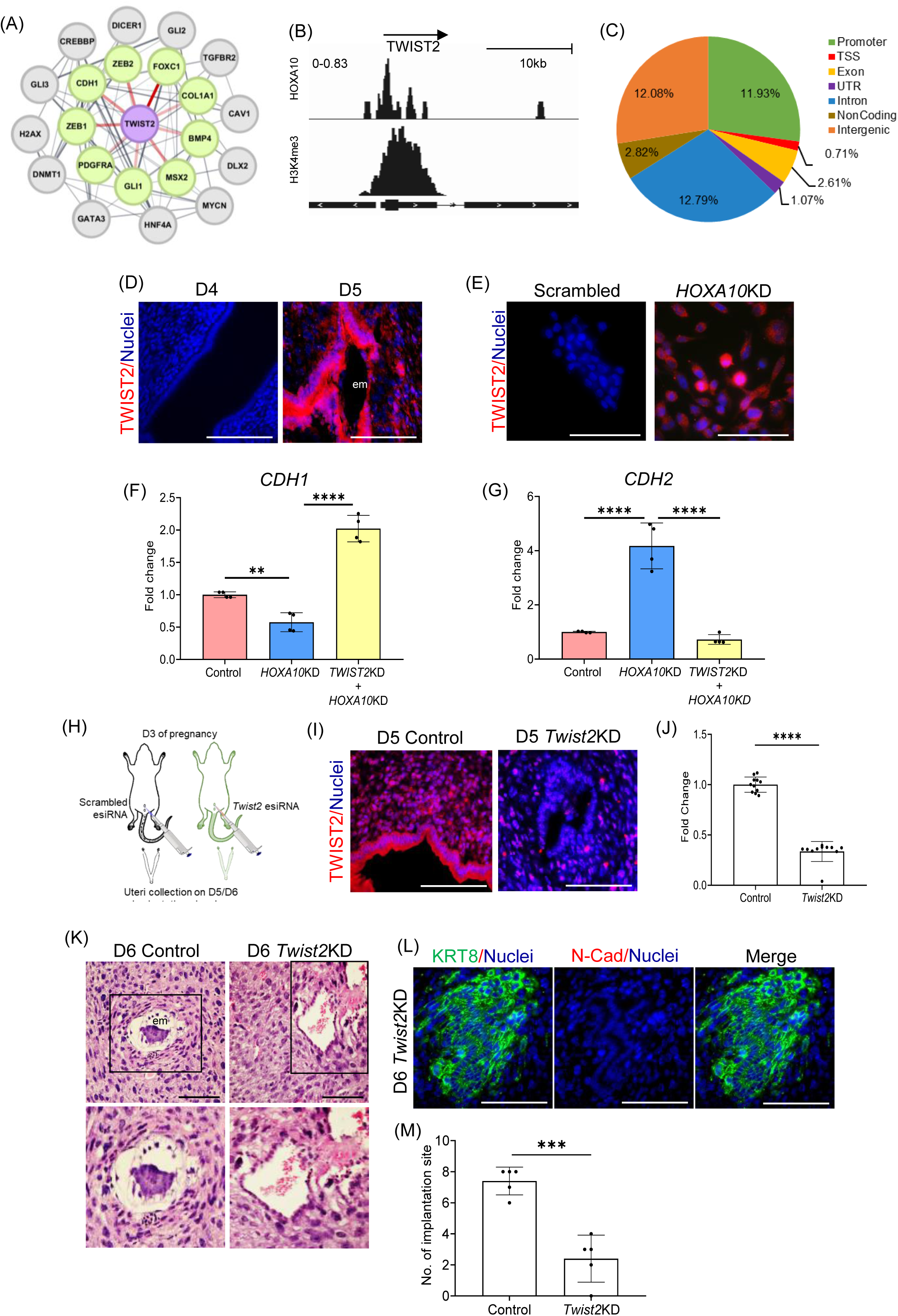
Mutual Antagonism Between HOXA10 and TWIST2 Controls pEMT and Initiates Implantation. **(A)** Gene regulatory PPI network depicts TWIST2 (purple node) interaction with various downstream target genes, the thickness of the edges represents the confidence of the interaction. **(B)** Occupancy of HOXA10 in *TWIST2* gene is shown in IGV plots, H3K4Me3 marks the promoters of the same genes. The arrow depicts the direction of gene transcription. **(C)** The pie chart illustrates the percentage of TWIST2 peaks located in different genomic regions of HOXA10 motifs. **(D)** Immunofluorescence for EMT transcription factors TWIST2 on D4 (09:00h) and D5 (09:00h) in the mouse uterus. **(E)** Immunostaining for TWIST2 in control and *HOXA10*KD endometrial epithelial cells. **(F)** mRNA levels of *CDH1* (normalize to 18s) in Control, *HOXA10*KD and *TWIST2*KD. Y-axis is a fold change where values obtained from control cells were taken as 1. Data is the mean ± SD for the four independent replicates (n=4). **(G)** mRNA levels of *CDH2* (normalize to 18s) in Control, *HOXA10KD* and *TWIST2KD*. Y-axis is fold change where values obtained from control cells were taken as 1. Data is the mean ± SD for the four independent replicates (n=4). **(H)** Schematic representation of experimental set up. Mouse uteri of control and *Twist2* esiRNA (*Twist2*KD) treated animal were collected on D5/D6. **(I)** Immunofluorescence (on D5 09:00h) for TWIST2 in the uteri of mice treated with scramble esiRNA (control) and *Twist2* esiRNA (*Twist2*KD). **(J)** Quantification of immunostaining for TWIST2 in control and *Twist2*KD (n=3). **(K)** Haematoxylin and Eosin-stained sections of control and *Twist2*KD mouse uterus on D6. Boxed images are zoomed below (n=3). **(L)** Immunofluorescence (on D6) for KRT8 and N-Cad in the uteri of mice treated with Twist2 *esiRNA* (n=3). **(M)** Quantification of the number of implantation sites in control and *Twist2*KD (n=5). em = embryo, Scale bar = 100µm. **** is p<0.0001, *** is p<0.001 and ** is p<0.05with mean ±SD values are shown.

To explore this, we examined TWIST2 expression across the implantation window. On D4, when HOXA10 levels are high, TWIST2 was absent in luminal epithelial cells. By D5, as HOXA10 expression reduced, TWIST2 was strongly induced in cells within the implantation crypt (Fig. 8D). *In vitro*, TWIST2 expression was markedly elevated in *HOXA10*KD cells, localizing robustly to the nucleus (Fig. 8E), suggesting that HOXA10 normally acts to repress TWIST2 in the epithelial state.

To test whether TWIST2 activation is functionally required for the HOXA10-driven pEMT, we knocked down *TWIST2* in *HOXA10*KD cells. This rescue experiment showed that repression of *TWIST2* significantly restored *CDH1* expression and reduced *CDH2* levels (Fig. 8F and G), indicating that *TWIST2* acts downstream of HOXA10 to drive the mesenchymal program.

We next asked whether this mechanism operates *in vivo*. *Twist2* esiRNA was administered intrauterine on D3, and uterine tissues were analysed on D5/D6 (Fig. 8H). In control animals, *TWIST2* was robustly expressed in the luminal epithelium on D5, whereas *Twist2* knockdown significantly reduced its expression (Fig. 8I and J). Histologically, while the control uteri showed complete epithelial clearance by D6 with embryos embedded in the stroma, the *Twist2* knockdown group retained an intact epithelial layer, resembling that of the D5 state (Fig. 8K). Co-staining for KRT8 and N-Cadherin confirmed that these retained cells had high KRT8 and lacked N-Cadherin, indicative of their epithelial nature and consistent with a blockade of pEMT (Fig. 8L).

Finally, we quantified implantation success in the absence of TWIST2. Compared to scrambled controls, *Twist2* knockdown significantly reduced the number of implantation sites (Fig. 8M). These findings underscore the requirement of TWIST2 activation for epithelial remodelling and successful embryo implantation.

## Discussion

Viviparity has driven eutherian mammals to evolve distinct strategies for embryo implantation into the endometrial stroma. In this study, we identify a novel strategy where mutually antagonistic regulatory circuit involving the transcription factors HOXA10 and TWIST2 governs pEMT in the endometrium to facilitate embryo implantation and establish pregnancy. Embryo implantation begins with the attachment of a competent blastocyst to the luminal epithelium of the receptive endometrium, followed by breaching of this epithelial barrier and invasion into the underlying stroma (Ashary *et al*, 2018). While multiple forms of cell death have been proposed to mediate epithelial cell displacement (Parr *et al*, 1987; Li *et al*, 2015; Akaeda *et al*, 2021), some evidence suggests the occurrence of EMT in the endometrial epithelial cells at the time of implantation (Uchida *et al*, 2012; Denker, 2016; Gou *et al*, 2019; Tiwari *et al*, 2021; Cheng *et al*, 2021). However, whether these endometrial epithelial cells undergo complete transition to a mesenchymal phenotype or acquire a pEMT state is not known and molecular mechanisms that drive EMT in the endometrial epithelium for embryo implantation are unknown. Here, we provide the first convincing demonstration that embryo implantation involves a form of pEMT, where luminal epithelial cells acquire migratory and mesenchymal traits without fully losing epithelial identity. Unlike pathological EMT, which is often sustained, the pEMT observed during implantation is spatially confined and tightly regulated, the hallmarks of physiological epithelial plasticity. Our findings not only resolve long-standing uncertainties about the nature of epithelial remodelling during implantation but also position the endometrium as a powerful *in vivo* model to study the transcriptional control of pEMT in adult tissues.

HOXA10 is a transcription factor critical for uterine development and is indispensable for fertility (Mishra & Modi, 2024). Although it is essential for the acquisition of endometrial receptivity, we and others have shown that its expression is specifically reduced in the luminal epithelium at the time of implantation in monkeys and canines (Godbole *et al*, 2007; Guo *et al*, 2009). We extend this observation to rodents, demonstrating significant HOXA10 downregulation in the luminal epithelium of both mouse and hamster uteri at implantation sites, suggesting an evolutionarily conserved phenomenon. This pattern is consistent with its regulation by progesterone signalling, as HOXA10 is a progesterone receptor (PGR)-responsive gene (Bhurke *et al*, 2016; Gebril *et al*, 2020), and PGR itself is downregulated in the luminal epithelium at implantation (Tan *et al*, 1999). Despite this correlation, the functional significance of HOXA10 downregulation during implantation has remained elusive.

To investigate the functional role of HOXA10, we silenced its expression in human endometrial epithelial cells and performed RNA-seq. HOXA10 loss led to altered expression of genes associated with tissue migration, actin cytoskeletal remodelling and focal adhesion dynamics, the hallmarks for cellular phenotype for motility. Live-cell imaging confirmed that loss of HOXA10 leads to pronounced migratory behaviour of the epithelial cells, suggesting that HOXA10 acts to restrain epithelial cell motility. At the site of implantation, *in vitro* imaging and interpretations from sections of the implantation sites in primates suggested that the luminal epithelial cells characteristically move away from the apposed embryo to pave a way for it to implant (Enders *et al*, 1983; Enders, 2000; Uchida *et al*, 2012; Siriwardena & Boroviak, 2022). To test whether HOXA10 loss enables this directional migration, we co-cultured trophoblast spheroids with *HOXA10*KD cells and observed outward epithelial cell migration from the site of spheroid attachment, creating a distinct clearance zone. While the experimental constraints do not allow us to observe this phenomenon *in vivo*, we observed premature epithelial sloughing in the implantation crypts of *Hoxa*10 hypomorphic mice, implying that HOXA10 downregulation promotes epithelial cell displacement at the implantation site.

To understand how HOXA10 regulates epithelial cell migration, we mapped its genome-wide occupancy and intersected this data with the transcriptome of *HOXA10*KD cells. We identified 1,246 differentially expressed direct target genes, many of which were associated with EMT and tissue migration pathways. Further, nearly 36% of EMT-related genes curated from the EMTome and dbEMT databases were found to be dysregulated in *HOXA10*KD cells, with most epithelial genes being downregulated and mesenchymal genes upregulated. These findings indicate that HOXA10 regulates cell migration in part by modulating EMT programs. We also found that *HOXA10*KD cells co-expressed E-Cadherin and N-Cadherin with variable intensities and exhibited elevated pEMT transcriptomic scores, consistent with a hybrid phenotype. *In vivo*, endometrial epithelial cells in *Hoxa10* hypomorphs similarly co-expressed E-Cadherin and N-Cadherin, confirming that HOXA10 loss is sufficient to induce pEMT in the uterine epithelium. The acquisition of the pEMT phenotype in the luminal epithelium at this conjuncture is conceivable as pEMT cells have higher collective migratory properties (Saxena *et al*, 2020), which are required for the luminal folding and formation of the implantation crypt. Such dynamic folding of the epithelium would require extensive migration of the epithelial sheets and EMT would certainly be required for such an activity. Indeed, altered LE folding and disrupted expression of EMT genes are observed in LE cells of mice knockout for planar cell polarity-related genes (Roberts *et al*, 2025). pEMT would also aid in loosening the tight epithelial cell-cell interactions and make way for the trophoblast cells to penetrate between them and initiate breaching.

While EMT is governed by a core set of transcription factors, both mathematical models and experimental data suggest that cells can adopt multiple partial and stable epithelial/mesenchymal (E/M) states due to dynamic interactions within EMT gene regulatory networks (McDermott *et al*, 2025). Perturbations in these networks by key transcription factors can tip the balance, giving rise to pEMT phenotypes (Rashid *et al*, 2022; Jain *et al*, 2024). We next asked whether HOXA10 loss perturbs such networks to drive epithelial cells toward a mesenchymal fate. Indeed, PPI network analysis of HOXA10 direct targets revealed two interconnected gene clusters, one associated with epithelial functions and the other with mesenchymal programs, and both clusters were enriched for processes linked to cell migration and motility. Consistently, we observed that HOXA10 occupancy extended across promoters of several epithelial genes (downregulated upon HOXA10 loss) and mesenchymal genes (upregulated upon HOXA10 loss), suggesting a dual regulatory role. These findings support a model wherein high HOXA10 expression during the receptive phase promotes epithelial identity by activating epithelial networks and repressing mesenchymal ones. Its downregulation at implantation shifts the balance, suppressing epithelial and enhancing mesenchymal gene expression to induce pEMT. Indeed, single-cell RNA-seq analysis of the luminal epithelial cells from the implantation stage corroborated this model, showing reduced expression of HOXA10-driven epithelial genes and increased expression of mesenchymal targets. Thus, HOXA10 functions as a core component of the epithelial "team" a transcriptional network that maintains epithelial state. Although few transcription factors are known to perform this stabilizing role, our findings identify HOXA10 as a key player in this process. Intriguingly, we also found enrichment of KLF-family transcription factor motifs in the HOXA10 cistrome; the KLF family of transcription factors have known roles in mesenchymal to epithelial transition (Subbalakshmi *et al*, 2021; Wang *et al*, 2023b), suggesting possible cooperative regulation between HOXA10 with other known epithelial-state inducers.

EMT is driven by a core set of master regulators of the SNAI, ZEB, and TWIST families, which coordinate the cellular reprogramming from an epithelial to a mesenchymal state (Huang *et al*, 2022). Within the PPI network of HOXA10-regulated genes, we identified ZEB1, ZEB2, and TWIST2. Of these, TWIST2 was identified as a direct HOXA10 target, with binding motifs in its promoter. In the mouse, we observed that TWIST2 is absent from the luminal epithelium prior to implantation but is strongly induced within the implantation crypt, coinciding with HOXA10 downregulation. Similarly, *HOXA10*KD cells showed upregulated TWIST2 expression, confirming that HOXA10 normally represses this mesenchymal transcription factor. Although this may appear to be a straightforward genetic switch, modelling studies have suggested that EMT and pEMT arise from mutually antagonistic transcriptional “teams.” In such networks, members of one module repress those of the opposing module, and the relative abundance of key players determines cell fate (Hari *et al*, 2022; Sahoo *et al*, 2025). Based on this framework, we hypothesized a mutual antagonism between HOXA10 and TWIST2. Indeed, knocking down TWIST2 in *HOXA10*KD cells restored *CDH1* and suppressed *CDH2* expression, indicating reversion toward an epithelial phenotype. *In vivo*, loss of TWIST2 impaired epithelial clearance at implantation sites; epithelial cells retained their identity, and the crypt structure was disorganized, resulting in implantation failure. Thus, TWIST2 activation is both necessary and sufficient to drive epithelial remodelling through pEMT and enable embryo implantation. These data support the view that pEMT involves the coordination of tightly regulated sub-networks where disruption of HOXA10 relieves repression of TWIST2-driven networks and initiates pEMT, and suppression of TWIST2 reverts the cells to their epithelial state.

Together, our findings uncover a conserved transcriptional circuit involving HOXA10 and TWIST2 that orchestrates a pEMT program in the endometrial luminal epithelium to enable successful embryo implantation. By acting as a stabilizer of epithelial identity, HOXA10 represses mesenchymal drivers such as TWIST2; its downregulation allows for a reversible, hybrid epithelial-mesenchymal state that facilitates epithelial remodelling without full mesenchymal transformation. This mechanism reveals how precise and transient modulation of epithelial plasticity supports tissue restructuring in response to developmental cues. Given the parallels between implantation and other physiological or pathological processes involving regulated EMT, such as wound healing, fibrosis, and cancer metastasis, our work highlights HOXA10 as a potential node in broader epithelial-mesenchymal regulatory networks. These insights open new avenues to explore how tissue plasticity is governed during dynamic epithelial transitions across organ systems.

## Summary

Collectively, our results uncover a previously unrecognized gene regulatory switch involving HOXA10 and TWIST2 that governs epithelial plasticity during implantation (Fig. 9). HOXA10 maintains epithelial identity by activating epithelial genes and repressing mesenchymal programs, including TWIST2. Its downregulation at the implantation site permits TWIST2 induction, triggering pEMT and enabling epithelial remodelling necessary for embryo invasion. This mutual antagonism provides a mechanistic framework for understanding how the uterine epithelium transitions into a permissive state for implantation.

**Fig. 9:**
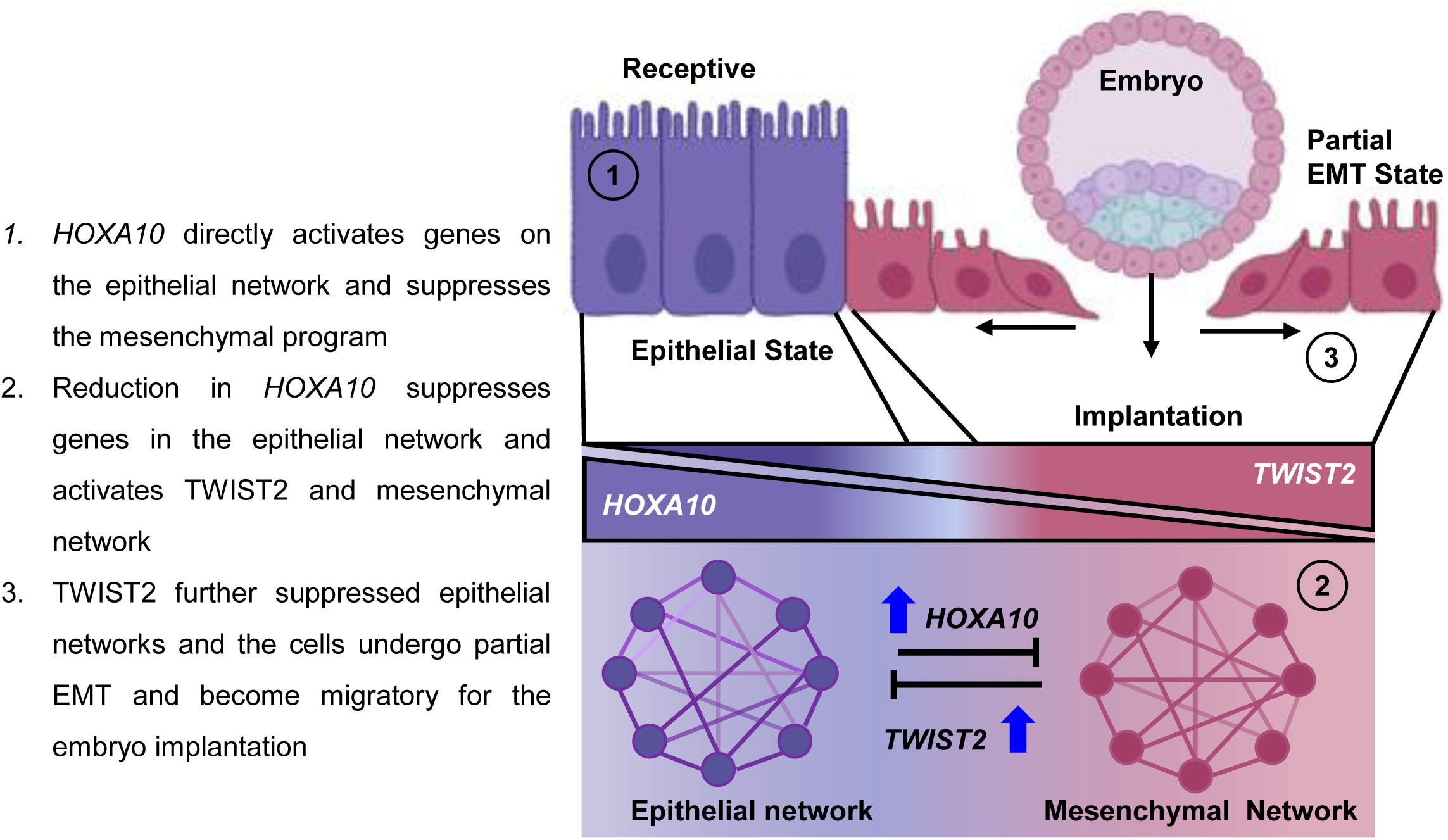
Summary illustrating the Process of Endometrial Epithelial Breaching for Initiation of Pregnancy.

## Materials and Methods

### Ethics Statement and Animal Tissue Collection

The use of animals was approved by the Institutional Animal Ethical Committee of ICMR-NIRRCH (project numbers 18/17 and 04/22) and Banaras Hindu University (BHU/DoZ/IAEC/2018-19/035). Adult (6–8-weeks old) C57BL/6 female mice housed in the experimental animal facility of ICMR-NIRRCH were mated with males of proven fertility and the day of vaginal plug was designated as day 1 of pregnancy. Uteri were collected on Day 4 (D4 09:00h), Day 5 morning (D5 09:00h), Day 5 evening (D5 21:00h) and Day 5 midnight (D5 24:00h). These time points were chosen as they span the period of receptivity, apposition, and implantation in mice (Li *et al*, 2015). HOXA10 hypomorphs are mice transgenic for a *Hoxa10* shRNA and have reduced expression of HOXA10 in their uteri as compared to their wild-type counterparts. The development and characterization of these mice are detailed elsewhere (Mishra *et al*, 2022). These animals were housed at the Experimental Animal Facility of ICMR-NIRRCH and the uteri were collected from either non-pregnant or on D5 (09:00h) of transgenic animals. The uteri from non-transgenic littermates were used as wild-type controls. Adult female golden hamsters (*Mesocricetus auratus)* housed in the animal house facility of BHU were mated with fertile males and the day of detection of sperm in a vaginal smear was considered as day 1 of pregnancy. Uteri from these animals were collected on day 5 (D5) which coincides with the day of implantation in this species (Reese *et al*, 2008). HOXA10 stained endometrium sections of bonnet monkey described previously (Godbole *et al*, 2007). were used.

### Histology and Immunofluorescence

Uteri at designated time points were collected and fixed in 4% paraformaldehyde (PFA, Sigma-Aldrich), paraffin-embedded and sectioned. The sections were stained using Hematoxylin and Eosin as detailed earlier (Mishra *et al*, 2020). Multiplex immunofluorescence was performed on paraffin sections using a commercial kit (PerkinElmer or Biotium), as described previously (Singh *et al*, 2023). The details of the antibodies used in the study and their optimized dilutions are given in (Table S3). Briefly, uterine sections were deparaffinized, and blocked in 2% Bovine Serum Albumin (BSA, Sigma-Aldrich). The antigenic sites were unmasked by incubating the sections in Tris-EDTA buffer (10 mM, pH 9) at 90°C for 30 min and probed with primary antibody. Post-washing, the sections were incubated with HRP conjugated secondary antibody, followed by tyramide signal amplification (TSA)-conjugated fluorophores for signal detection. For multiplexing, after the first round of probing, the sections were incubated for 30 min at 90 °C in the Tris-EDTA buffer and probed with the next antibody. The sections were counterstained with DAPI (Sigma-Aldrich) and mounted with EverBrite™ Hardset Mounting Medium (Biotium). For immunofluorescence on cultured cells, they were trypsinized and seeded on sterilized coverslips and cultured till the desired confluency. Cells were fixed in chilled 70% methanol, washed, blocked in BSA and then incubated with primary antibody overnight at 4°C. Next day, after washing, the cells were incubated with fluorescently conjugated secondary antibody (Invitrogen), and the nuclei were counterstained with DAPI and mounted as above.

Imaging was done on an inverted fluorescence microscope equipped with sCMOS camera (Leica Microsystems DMi8, Germany). The images were analyzed, and fluorescence intensities from selected areas were quantified in Leica Microsystems using LAS X software. Independent areas of luminal epithelium in non-consecutive sections were selected using the Region of Interest (ROI) function and the intensities of the desired signals were estimated. For estimating co-expression, the multi-colored images were imported in ImageJ (Fiji) and JACoP Co-localization plugin was used to determine the Pearson’s correlation coefficient in the luminal epithelial cells. For single-cell quantification, each luminal epithelium cell surrounding the embryo was individually selected using the ROI function, and the fluorescence intensities were estimated using LAS X software. At least 50 non-overlapping cells per implantation site were quantified. Similarly single cell quantification of endometrial epithelial cells was estimated using LAS X software.

### HOXA10 Knockdown in the Endometrial Epithelial Cells

RL95-2 cell line (henceforth referred to as RL95) are human endometrial epithelial cells and highly adhesive to trophoblast spheroids and are considered a suitable model to investigate human embryo implantation *in vitro* (Hannan *et al*, 2010). RL95 cells (ATCC-CRL-1671) were cultured on gelatin-coated plates in DMEM-F12 (Gibco) and 10% fetal bovine serum (FBS, Gibco). To silence HOXA10, cells were transfected with a plasmid containing a *HOXA10* shRNA (V3LHS_300885; Dharmacon) or a scrambled shRNA (V3LHS_65890S; Dharmacon) using X-tremeGENE™ HP DNA Transfection Reagent (Roche). Both these plasmids have been prevalidated *in vitro* and *in vivo* (Mishra *et al*, 2022). After 48h of transfection, the cells were challenged with 2ug/ml of puromycin (Gibco) for 48h, and the surviving cells were sub-cultured and used for further experiments. The knockdown efficiency of HOXA10 was examined by qPCR (material and methods S2 and Fig. S3). Cells were regularly tested for mycoplasma using PCR (HiMedia).

### RNAseq

RNA was isolated from 70% confluent *HOXA10*KD and control endometrial epithelial cells using the RNEasy mini kit. High-quality RNA (RIN values >9) was subjected to standard paired-end RNA-Seq library preparation using NEBNext® Ultra™ II Directional RNA Library Prep Kit (NEB-E7760S). Sequencing was performed on an Illumina NovaSeq 6000 platform at Bionivid Technology (Bangalore, India). Low-quality reads were trimmed and the raw sequence data was processed, and aligned to the human reference genome (hg38). Raw counts were calculated using Feature Counts and differential gene expression (DEGs) analysis was performed using EdgeR. Genes with a Log2 fold change ±2 and a P-adjusted value of 0.05 were considered as differentially expressed.

### Time-Lapse Microscopy for Cell Migration

Cells knockdown for HOXA10 (*HOXA10*KD) and scrambled (control) were seeded into 96-well microplates (Revvity) and cultured in DMEM-F12 medium containing 10% FBS and 2ug/ml puromycin. The next day, cells were imaged every 30 min for 24 h using the Operetta CLS High-Content Analysis system equipped with 37 °C, 5% CO2 environmentally controlled chamber (Revvity). Chemotaxis and migration plugins (Harmony High-Content Imaging and Analysis Software) were used to estimate cell migration and displacement.

### Spheroid Induced Cell Displacement Assay

To test if embryonic spheroids have an effect on the displacement and migration of endometrial epithelial cells, spheroid induced cell displacement assay was performed as described in (Iwanicki *et al*, 2011), with minor modifications. Briefly, mycoplasma free trophoblast cells (HTR8/SV-neo cells, ATCC-CRL-3271) cultured in DMEM-F12 and 10% FBS were seeded in low attachment plates (PerkimElmer) and 24h later the spheroids were labeled with CellTracker™ Red CMTPX (Invitrogen). Approximately 5 spheroids per well were co-cultured with monolayers of CellTracker™ Green CMFDA (Invitrogen) labeled control or *HOXA10*KD cells. After 24h the wells were imaged under a fluorescence microscope (Leica Microsystems DMi8).

For live cell imaging, the *HOXA10*KD cells were seeded into 96-well microplates (PerkinElmer) and grown until 90% confluency. Trophoblast spheroids were allowed to attach to these cells for 2h and the endometrial cells in the area of the spheroids were imaged every 30 min for 12h in the Operetta CLS High-Content screening system. The cell migration patterns were analyzed using Harmony High-Content Imaging and Analysis Software.

### Identification of HOXA10 Occupancy in Endometrial Epithelial Cells using CUT&RUN

Mycoplasma free endometrial epithelial cells (RL95 cells) were treated with estrogen (10^-8^M) and progesterone (10^-6^M) (both Sigma-Aldrich) for 48h. Preliminary experiments had shown that at these concentration HOXA10 expression is robustly increased in RL95 cells. The cells were collected by trypsinization and subjected to CUT&RUN using a commercial kit (CUT&RUN Assay Kit #86652 Cell Signaling Technology). Briefly, 100,000 cells were lysed and nuclei were washed thrice and incubated overnight at 4°C with a validated HOXA10 antibody (50ug). The validation of this antibody is provided in (Fig. S6). Excess primary antibody was removed by multiple washings and the nuclei were incubated with a pAG-MNase. It was activated with CaCl_2_ at 4°C to initiate DNA digestion for 30 min and then stopped using the stop buffer. The released DNA fragments were purified using the conventional phenol-chloroform DNA extraction method. DNA library preparation was done as detailed (Skene & Henikoff, 2017) and sequencing on NovaSeq 6000 platform was outsourced (Clevergene, Bangalore, India). High-quality reads were aligned to the human reference genome (hg38), and peak calling was performed using SEACR, with the green list employed as a control (de Mello *et al*, 2024). The top 10,000 peaks were annotated, and motif enrichment analysis was conducted using the MEME (Multiple Em for Motif Elicitation) suite version 5.5.7.

### RNAseq and CUT&RUN Data Analysis and Visualization

Gene enrichment analysis of DEGs was performed using WebGestalt (Elizarraras *et al*, 2024), and pathways of interest were visualized using KEGG Mapper (Xie *et al*, 2011). Data visualization included Integrative Genomics Viewer (IGV), heatmaps, volcano plots, Sankey plots, alluvial plots, and bubble plots. These visualizations were generated using the ggplot2 and SRPlot packages within the R statistical environment. Protein-protein interaction (PPI) networks were constructed using the Cytoscape StringApp. HOXA10 co-enriched motifs were analyzed MEME-FIMO suite version 5.5.7. The single-sample gene set enrichment analysis (ssGSEA) algorithm (Barbie *et al*, 2009), as implemented in GSEAPy (Fang *et al*, 2023), was used to calculate ssGSEA scores for the partial EMT (pEMT) signature (Puram *et al*, 2017). Pearson correlations between gene expression levels and either HOXA10 expression or pEMT scores were computed using the Pearson function from the SciPy library (Virtanen *et al*, 2020).

### Single cell RNA-seq data analysis

To study the expression of epithelial and mesenchymal gene in mouse luminal epithelium at single-cell resolution, we analyzed scRNA-seq data [GSA: CRA011267] of mouse uteri on day 3 and day 5 of pregnancy (Wang *et al*, 2023a). Gene expressed in the *Tacstd2* (marker for luminal epithelial cell) (Wang *et al*, 2023a) were extracted and analysed.

### Knockdown of Twist2 in vitro and in vivo

To silence TWIST2 *in vitro*, cells were transfected with a 75pmoles *TWIST2* esiRNA (Sigma-Aldrich) using Lipofectamine 3000 (Invitrogen) reagent. After 48h of transfection, the cells were sub-cultured and used for further experiments.

To knockdown TWIST2 in the uterine cells *in vivo*, 200ng of *Twist2* esiRNA or scrambled esiRNA was incubated in Lipofectamine 3000 reagent and 20 µl of the complex was delivered intraluminally into the uterus of day 3 (D3) pregnant mice using the NSET™ device (ParaTechs) as per the manufacturer’s instructions. Briefly, the NSET device was attached to a pipette and the large speculum was used to hold open the vagina and visualize the cervix. The NSET tip was inserted up to the utero-cervical junction and the solution was expelled. After 48 or 72h of administration (D5 or D6 of pregnancy) the uteri were collected, fixed, and processed as above for serial sectioning. The numbers of implantations per animal were estimated in uterine sections by counting the number of implantation crypts (on D5) or embryos embedded in the uterine stroma (on D6). For each animal, every 5th slide was assessed and at least 10 slides/animal were evaluated. The details of *Twist2* esiRNA is given in (Table S4). The efficiency of knockdown in the uterine sections was determined using immunofluorescence for TWIST2.

### Statistics

The mean ± SD for all the experimental data was calculated and statistical analysis was done by Student’s t-test and one way ANOVA using GraphPad Prism, version 8. p<0.05 was accepted as statistically significant.

## Data Availability

All data generated in this study is available at European Nucleotide Archive (ENA) with Accession Number PRJEB82872 (RNA-seq data) and PRJEB83114 (CUT&RUN data).

## Author Contributions

DM and NA designed the research. NA, SS, S Shyamal, PN an HBV performed research. AM, AA, SH, MKJ contributed new reagents/analytic tools. DM, NA, SS, AB, S Shyamal, S Patil, HBV and MKJ analysed data. All author’s contributed to manuscript writing.

## Acknowledgement

We are grateful to Varun Suresh for valuable technical inputs, we are thankful to Dr Geetanjali Sachdeva (ICMR-NIRRCH) for kind gift of RL95-2 cells, we are thankful to Dr Bhakti Pathak (ICMR-NIRRCH) for inputs in establishing stable clones. ChatGPT-4.0 was used to assist with language editing of the manuscript, under the authors’ supervision.

## Funding

Department of Science and Technology - Science and Engineering Research Board (DST No: CRG/2018/002314) (DM)

Department of Biotechnology (BT/PR51181/MED/97/675/2023) (DM)

Indian Council of Medical Research (ICMR), Govt. of India (DM).

ICMR-Senior Research Fellowship (NA and AM)

Department of Biotechnology-BioCARe for funding (BT/PR50903/BIC/101/1307/2023) (S Shyamal)

Institution of Eminence, Banaras Hindu University (IoE-BHU), PDF (6031) (SH) Prime Minister’s Research Fellowship (PMRF) (HBV)

Param Hansa philanthropies (MKJ)

The manuscript bears the NIRRCH ID: RA/1817/01-2025.

