## Supplementary data for "HOXA10-TWIST2 Antagonism Drives Partial Epithelial-to-Mesenchymal transition for Embryo Implantation"

Nancy Ashary *et al.*

**This PDF file includes:**

Materials and Methods S1and S2
Figures S1 to S9
Tables S1 to S4
Legends for Movies S1 to S3
Supplementary References

### Materials and Methods

#### S1. Immunohistochemistry

Immunohistochemistry was performed as described (Mishra et al, 2022); Briefly, paraffin embedded sections were deparaffinized in xylene and rehydrated in grades of alcohol. Antigen was unmasked using Tris EDTA buffer (10 mM, pH 9) at 90 °C. The sections were blocked using 1% donkey serum (Jackson Immunology) for 1h. Further sections were probed with a primary antibody (HOXA10) for overnight. Next day sections were washed and incubated with a biotinylated secondary antibody followed by streptavidin-HRP (ABC kit Santa Cruz Biotechnology). 3, 3′-diaminobenzidine (DAB) (Sigma -Aldrich) was used as a chromogen and hydrogen peroxidase as a substrate for the detection. Sections were counterstained with haematoxylin and mounted in DPX. Slides were viewed under a bright field microscope (Olympus) and representative areas were photographed. "Fiji" version of ImageJ software was used to quantification of HOXA10 expression

.

#### S2. RT-PCR

Total RNA from cells were extracted using Trizol reagent (Invitrogen) as described previously (Mishra et al, 2022); and reverse-transcribed to cDNA (cDNA reverse transcriptase kit, Applied Biosystems). Real time PCR was performed using the CFX-96 thermal cycler (Bio-Rad) using SYBR green chemistry (Bio-Rad). The annealing temperature was optimized for each gene. Gene expression was normalized to the levels of 18S, and fold change was calculated.

| Gene Name | Primer sequence<br>5’-3’ |
|---|---|
| *HOXA10* | 5'-GCCCCTTCCGAGAGCAGAAAA-3'<br>5'-AGGTGGAGCCTGCGGCTAATCTCTA-3' |
| *CDH1* (E-Cadherin) | 5'-GAACAGCACGTACACAGCCCT-3'<br>5'-GCAGAAGTGTCCCTGTTCCAG-3' |
| *CDH2* (N-Cadherin) | 5'-GACGGTTCGCCATCCAGAC-3'<br>5'-TCGATTGGTTTGACCACGG-3' |

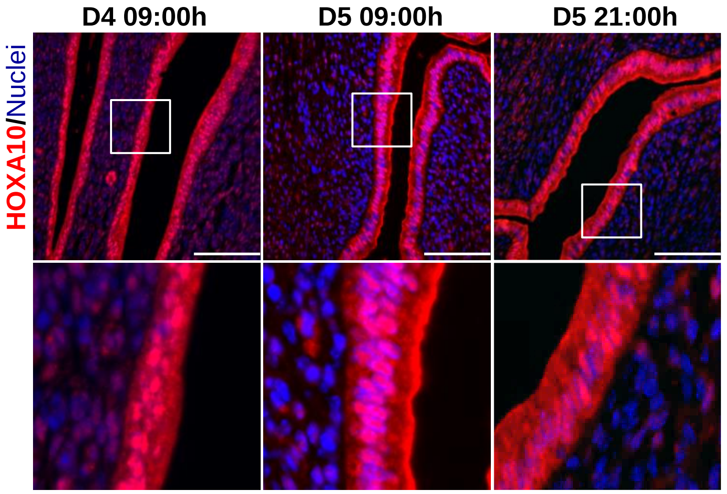


**Fig S1: Immuno-localization of HOXA10 at the inter-implantation site of mouse endometrium**
Selected area is boxed and zoomed in below image. Scale bar = 100µm, n=3

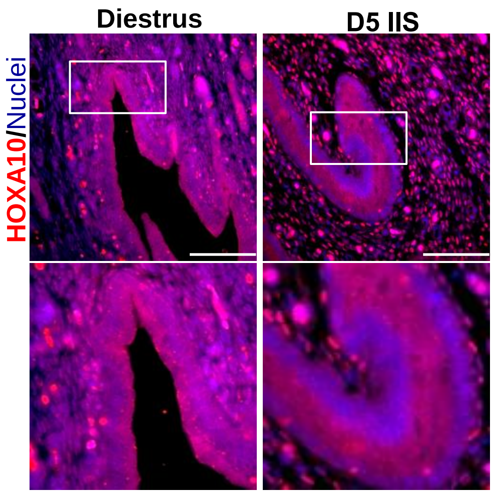


**Fig S2: Immuno-localization of HOXA10 in Diestrus stage and inter-implantation site (IIS) of Day 5 (D5) hamster endometrium**

Selected area is boxed and zoomed in below image. Scale bar = 100µm, n=3

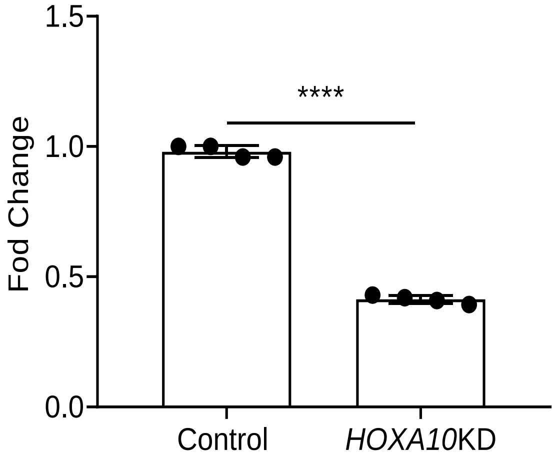


**Fig S3. Validation of *HOXA10* knockdown in endometrial epithelial cells (RL95-2)**
mRNA levels of *HOXA10* (normalize to 18s) in RL95-2 cells stably expressing scrambled shRNA (Control) and *HOXA10* shRNA (*HOXA10KD)*. Y-axis is fold change where values obtained from control cells was taken as 1. Data is the mean ± SD for the Four independent replicates. **** indicates significant difference as compared to control ($p < 0.0001$)

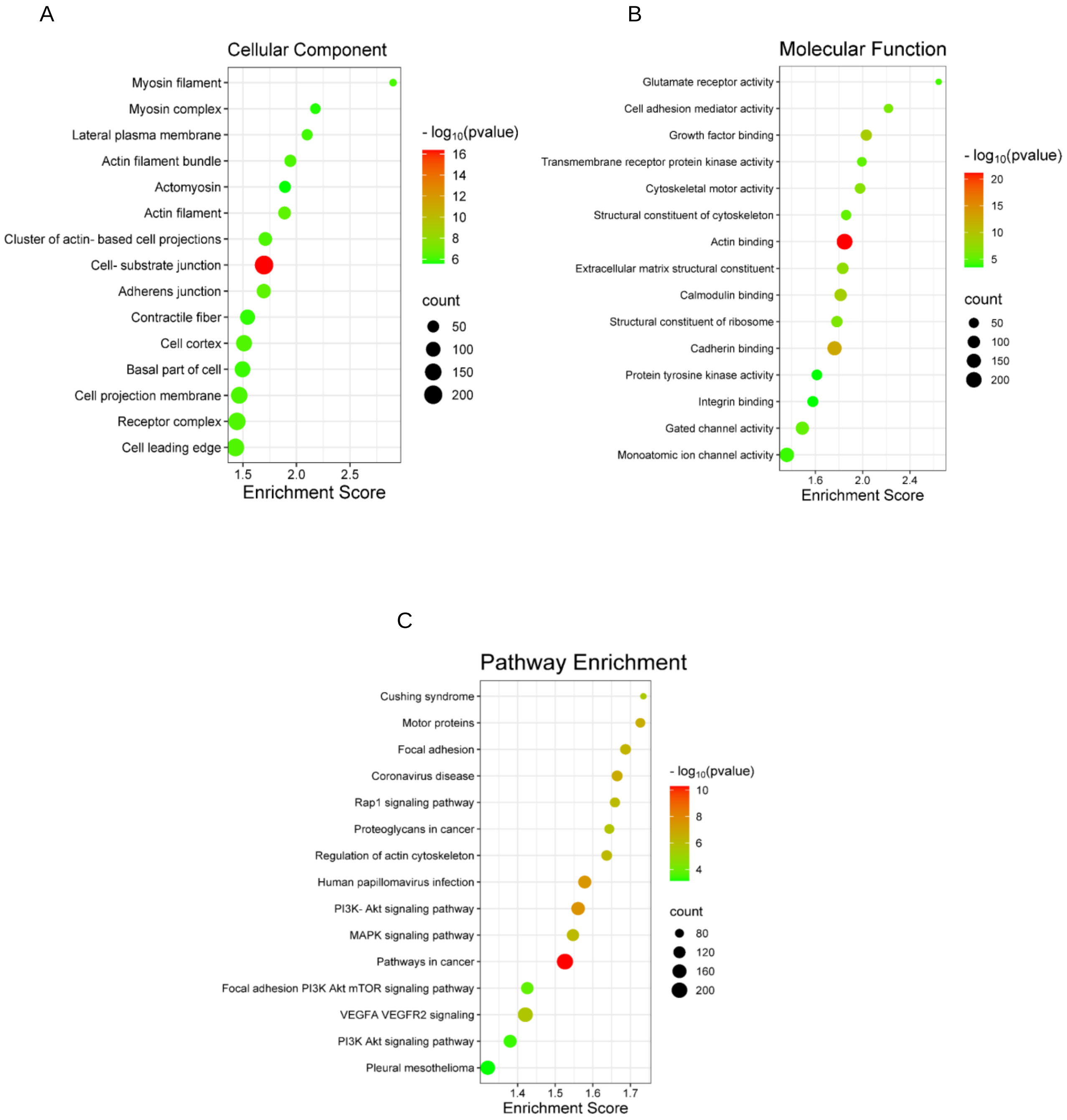


**Fig S4: Gene Ontology and Pathway Enrichment Analysis of the DEGs in the *HOXA10* knockdown in human endometrial epithelial cells (RL95-2)** This figure depicts the results of Gene Ontology (GO) enrichment analysis for **(A)** cellular component; **(B)** molecular functions, **(C)** pathways. Enrichment scores (x-axis) indicate the statistical significance of the enrichment for each term or pathway. Bubble size represents the number of genes associated with each term, and color indicates the -log10(p-value).

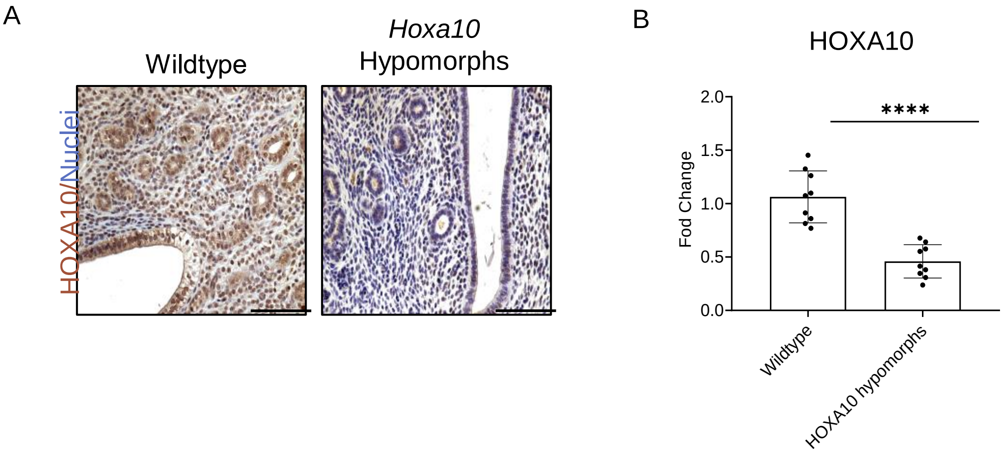


**Fig S5: Expression of HOXA10 in the uteri of Mice transgenic for shRNA against *Hoxa10* (*Hoxa10* hypomorphs). (A)** Immunohistochemistry of HOXA10 in wildtype and *Hoxa10* hypomorphs in diestrus stage mouse endometrium. **(B)** Graph represent intensity of HOXA10 immunostaining. Values on Y-axis are fold change where the mean value of controls was taken as 1. Scale bar = 100µm. ****in Graph represent statistically significant ($p<0.0001$) with mean ±SD values (n=3) are shown.

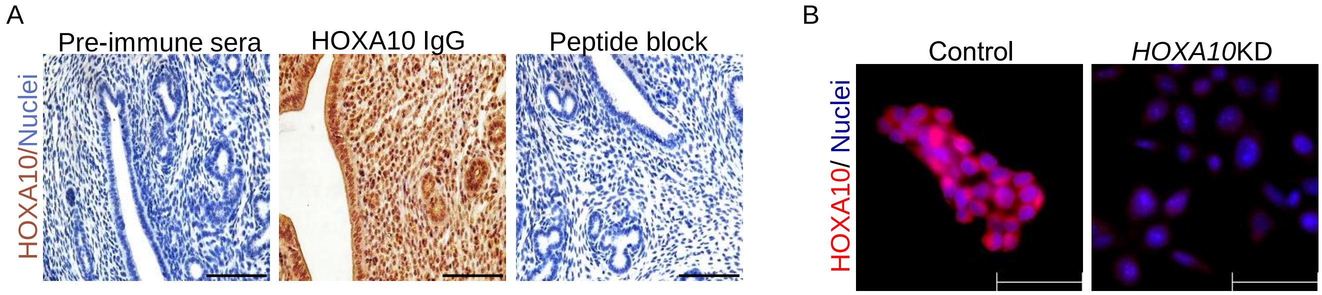


C CUT&RUN of HOXA10 in RL95-2 cells

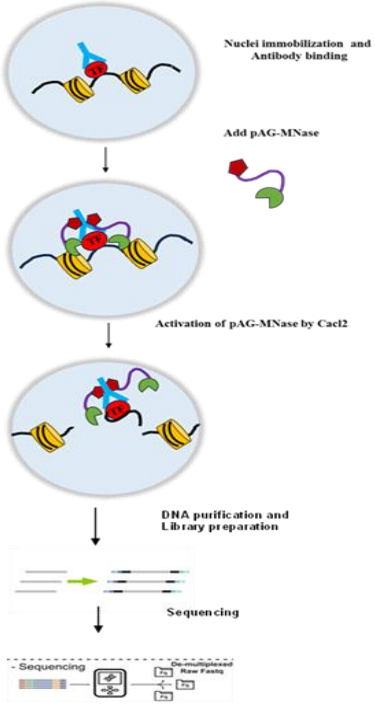


| HOXA10 Motif | pvalue |
|---|---|
| 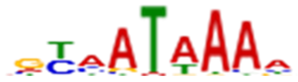 | 1.78e-3 |

**Fig S6: Validation of customized HOXA10 antibody.**

Rabbits were immunized with KLH conjugated 19mer peptide of HOXA10. Four boosters were given and the sera were collected and the IgG fraction was affinity purified . These experiments were done commercially (GenScript, USA). The generated antibody was validated as below.

**(A)** Immunohistochemistry in paraffin sections of diestrus stage mouse endometrium. Scale bar = 100µm. Section incubated with pre-immune sera/affinity purified IgG (HOXA10 IgG)/affinity purified IgG with excess of the peptide (peptide block) used for immunization. The results revealed intense staining in the sections incubated with the HOXA10 IgG but not in the pre-immune sera. This staining was not observed in section incubated with the peptide indicating its specificity. **(B)** Immunostaining for HOXA10 in scrambled shRNA (control) and *HOXA10* shRNA (*HOXA10*KD) endometrial epithelial cells (RL95-2). We also validated the antibody in human endometrial epithelial cells, where HOXA10 was knocked down using a validated shRNA (Mishra et al, 2022) and see fig S3 and S5). Intense signals for HOXA10 were detected in the nuclei of the control cells; these signals were reduced significantly in the *HOXA10*KD cells further demonstrating its specificity. Scale bar = 100µm. **(C)** To determine if the antibody identified the HOXA10 bound to the chromatin, CUT&RUN was performed using the affinity purified HOXA10 IgG. Nuclei of RL95-2 cells treated with Estrogen and Progesterone were incubated with the IgG and CUT&RUN was performed using a commercial kit as outlined in the figure and detailed in the methodology. HOXA10 motifs was significantly enriched within the top 5k high intensity peaks.

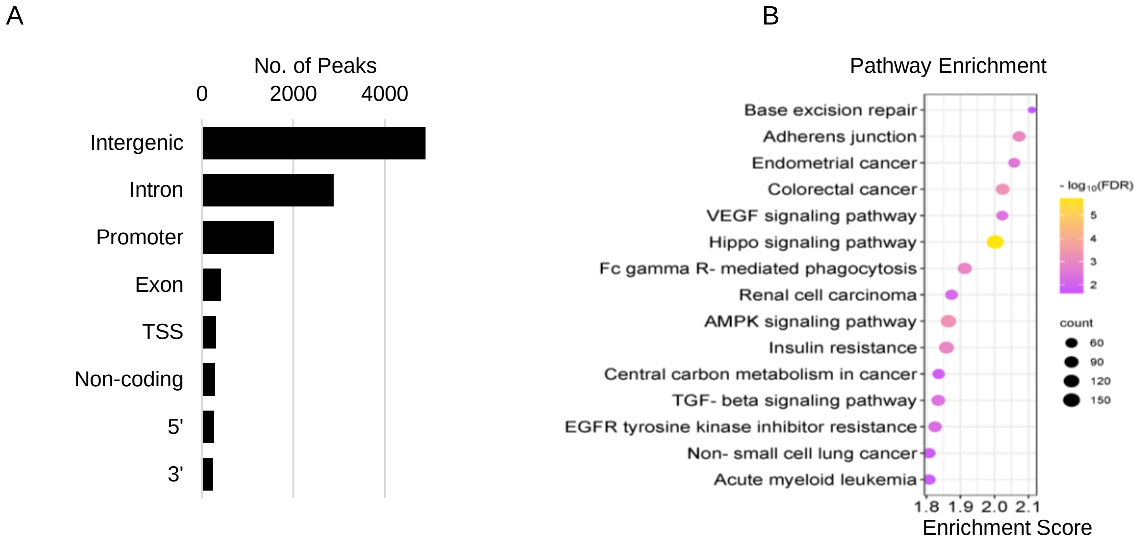


**Fig S7: Genomic localization of HOXA10 CUT&RUN peaks**

**(A)** Location statistics of top 10,000 high confidence peaks identified from CUR&RUN for HOXA10. **(B)** Pathways associated with the genes that have HOXA10 occupancy on their promoters and transcription start site (TSS).

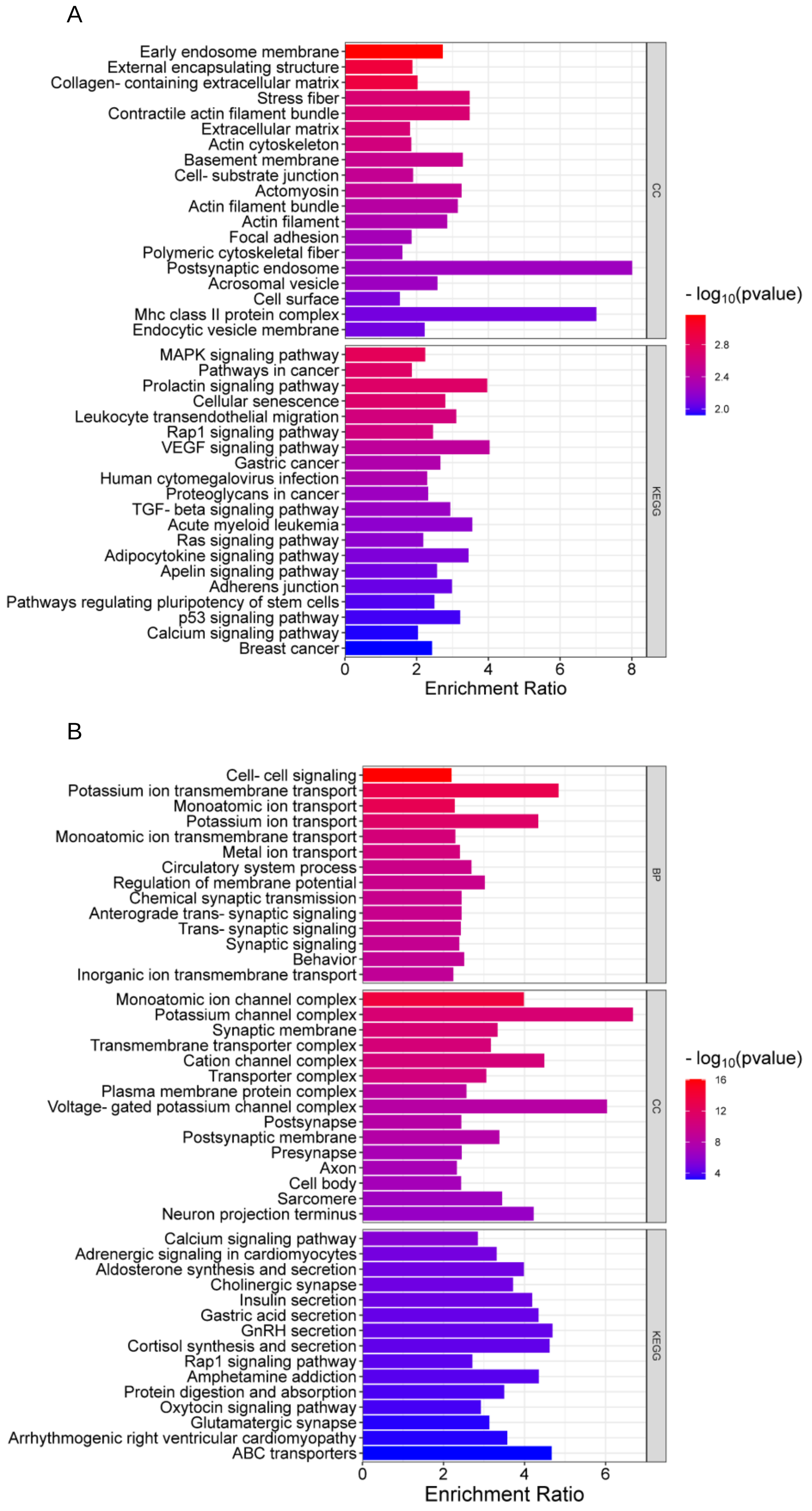


**Fig S8: Gene Ontology and Pathway Enrichment Analysis for Upregulated and Downregulated directs targets of HOXA10.**

Genes that were differentially expressed in *HOXA10*KD cells and whose promoters/TSS had HOXA10 occupancy (direct targets) were subjected to GSEA. Panel representing the GO terms for **(A)** upregulated direct targets and **(B)** down regulated direct targets

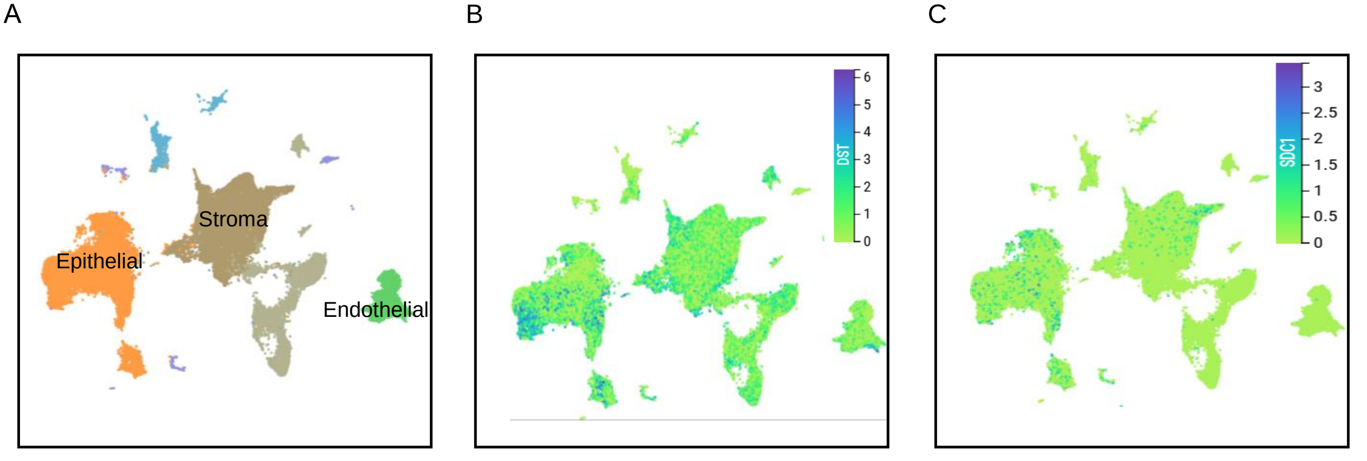


**Fig S9: Localisation of epithelial genes *DST* and *SDC1* in Human Endometrium**
Data for *DST* and *SDC1* was obtained from (https://www.reproductivecellatlas.org/endometrium-all). **(A)** UMAP representations colored by cell type of human endometrium. **(B)** UMAP representations of *DST* expression in different cell type of human endometrium. **(C)** UMAP representations of *SDC1* expression in different cell type of human endometrium

**Table S1. List of motifs co-enriched in the HOXA10 cistrome and have known roles in implantation.**

| | Role In Embryo Implantation |
|---|---|
| Estrogen receptor | Estrogen is a critical determinant that specifies the duration of the window of uterine receptivity for implantation (Ma *et al*, 2003 ,Okur et al, 2016). |
| KLF5 | Klf5 is critical to making the uterine luminal epithelium conducive to blastocyst implantation and growth.<br>Epithelium is retained in Klf5 deleted females past the WOI and Decidualization in impaired (Sun et al, 2012). |
| FOXA2 | FOXA2 is a critical regulator of uterine gland function, embryo implantation, and pregnancy establishment. Blastocyst attachment in adult FOXA2-deficient uteri is impaired due to stromal cell decidualization impairment. Hoxa10 expression was increased in the implantation sites of FOXA2 deleted mice (Kelleher et al, 2017). |
| KLF4 | KLF4 promotes decidualization of human endometrial stromal cells by regulating their autophagy level.<br>In adenomyotic human endometrial stromal cells decidualization id impaired, overexpression of KLF4 significantly reversed the decreased secretion of decidualization PRL (Mei et al, 2022). |
| KLF9 | BTEB1, the transcription factor of KLF9, by regulating Stromal progesterone receptor expression and transactivation, participates in the paracrine control of luminal epithelium proliferation by PGR and thus is important for establishment of a receptive uterus critical for successful implantation (Velarde et al, 2005). |
| | HOXA10-binding sites in the KLF9 promoter directly regulate KLF9 expression. Together they likely regulate progesterone action in endometrial epithelial cells (Du et al, 2010). |
| FOXO1 | Uterine ablation of Foxo1 using the progesterone receptor Cre (PgrCre) mouse model resulted in infertility due to altered epithelial cell polarity and apoptosis, preventing the embryo from penetrating the luminal epithelium. Foxo1 knocked Down mice indicated that endometrial FOXO1 was required for the temporal permeability of the LE (Vasquez et al, 2018). |
| SMADs | Uterine epithelial BMP/SMAD1/5 signalling is essential during early pregnancy and SMAD1/5 epithelial-specific deletion has detrimental effects on stromal cell decidualization and pregnancy development.<br>In Smad1/5 knocked down mice implantation failed with decrease in COX2 expression and FOXO cytoplasmic miss localization (Tang *et al*, 2022). |
| | Uteri from Smad1, smad5, smad4 and Amhr2 conditionally Knocked down females exhibit multiple defects in stroma, epithelium, and smooth muscle layers and fail to assemble a closed uterine lumen upon embryo implantation, with defective uterine decidualization that led to pregnancy loss at early to mid-gestation (Rodriguez *et al*, 2016). |
| | Inhibition of SMAD2/3 signalling disrupts organoid morphology, increases the glandular and secretory cell markers, FOXA2 and MUC1, and alters the genome-wide distribution of SMAD4.<br>TGFβ family signalling via SMAD2/3 controls signalling networks which are integral for endometrial cell regeneration and differentiation (Kriseman *et al*, 2023). |
| | TGFβ family signalling via SMAD2/3 controls signaling networks which are integral for endometrial cell regeneration and differentiation. Mechanistic studies in endometrial organoids show that inhibition of SMAD2/3 signalling disrupts organoid morphology, increases the glandular and secretory cell markers, FOXA2 and MUC1, and alters the genome-wide distribution of SMAD4 (Zhao *et al*, 2012). |
| | Inhibiting SMAD3 on day 3 of pregnancy in mice showed reduction in IGFBP-1 and decreased number of implanted embryos (Li *et al*, 2019). |
| STAT3 | STAT3 activation in luminal epithelium, regulates epithelial cell-cell junctions, polarity, and function during the receptive phase and also promotes stromal proliferation at the time of decidualization via paracrine growth regulatory signals originating in the epithelium (Pawar *et al*, 2013). |

**Table S2. List of Signaling pathways and its role in embryo implantation**

| Pathway | Role in embryo implantation |
|---|---|
| Hippo signaling | The study demonstrates the expressions of pYAP, YAP, TEAD1, and CTGF which are members of the Hippo signaling pathway, in the uterus of mice during the peri-implantation phase. The Hippo signaling pathway shows dynamic changes during the peri-implantation period and is involved in both implantation and decidualization. (Golal *et al*, 2023, Moldovan et al, 2025). |
| AMPK | AMPK activity is a key mediator of various steroid hormone-dependent processes during pregnancy establishment, including decidualization, uterine receptivity, and epithelial cell proliferation (Griffiths IV *et al*, 2020). |
| TGFBeta | Active TGF-B1, available from its latent complex, triggers SMAD3 which is crucial for embryo implantation. Loss of active TGF-B1 affects the development of blastocyst and endometrium receptivity that leads to the reduction in fetus number (Maurya *et al*, 2013). |
| | Expression levels of some immunomodulatory cytokines in endometrium are significantly increased even before the embryo invades the endometrium. The endometrial expression of TGFβ2, TGFβ2 receptor, PP14 and IL-6 were significantly up-regulated ($p < 0.05$) in pregnant animals as compared to non-pregnant animals, whereas the expression of LIF and its receptor remained unaltered in pregnant animals (Rosario *et al*, 2005). |
| VEGF | VEGF plays a crucial role in mediating the increase in estrogen-induced uterine vascular permeability, and it is essential for implantation (Rockwell *et al*, 2002). |

**Table S3. List of antibodies and their optimized dilutions used in this study**

| Primary Antibody | Company | Catalogue | Dilution |
|---|---|---|---|
| HOXA10 | GenScript. | customized | 1:50 |
| HOXA10 | Biomatik | CAE03449 | 1:200 |
| E-Cad | Abcam | ab15148 | 1:50 |
| N-Cad | Abcam | ab18203 | 1:100 |
| KRT8 | Abcam | ab154301 | 1:250 |
| TWIST2 | Novus Bio | NBP2-56209 | 1:100 |
| F-actin | Biotium | Phalloidin, CF®568 | 1:50 |
| Goat anti-rabbit IgG Alexa 568 | Invitrogen | A11011 | 1:1000 |

**Table S4. TWIST2 esiRNA and Control esiRNA details**

| esiRNA | Company | Catalogue |
|---|---|---|
| Mouse *Twist2* esiRNA | Sigma-Aldrich | EMU043621 |
| Human *TWIST2* esiRNA | Sigma-Aldrich | EHU227201 |
| Control esiRNA | Sigma-Aldrich | EHURLUC |

**Supplementary Movie legends**

**Movie S1:** Representative phase contrast timelapse of control endometrial epithelial cells (RL95). Cells were imaged every 30 min for 24h using the Operetta CLS High-Content Analysis system equipped with 37 °C, 5% CO2 environmentally controlled chamber (Revvity). Total elapsed time from first image is indicated at top left

**Movie S2:** Representative phase contrast timelapse of *HOXA10*KD endometrial epithelial cells (RL95). Cells were imaged every 30 min for 24h using the Operetta CLS High-Content Analysis system equipped with 37 °C, 5% CO2 environmentally controlled chamber (Revvity). Total elapsed time from first image is indicated at top left.

**Movie S3:** Representative phase contrast timelapse of *HOXA10*KD cells co-incubated with trophoblast spheroid (HRT8/SV-neo). Cells were imaged every 30 min for 12h using the Operetta CLS High-Content Analysis system equipped with 37 °C, 5% CO2 environmentally controlled chamber (Revvity). Total elapsed time from first image is indicated at top left.
