## Supplementary figures and images for "HOXA10-TWIST2 Antagonism Drives Partial Epithelial-to-Mesenchymal transition for Embryo Implantation"

### Movie S1

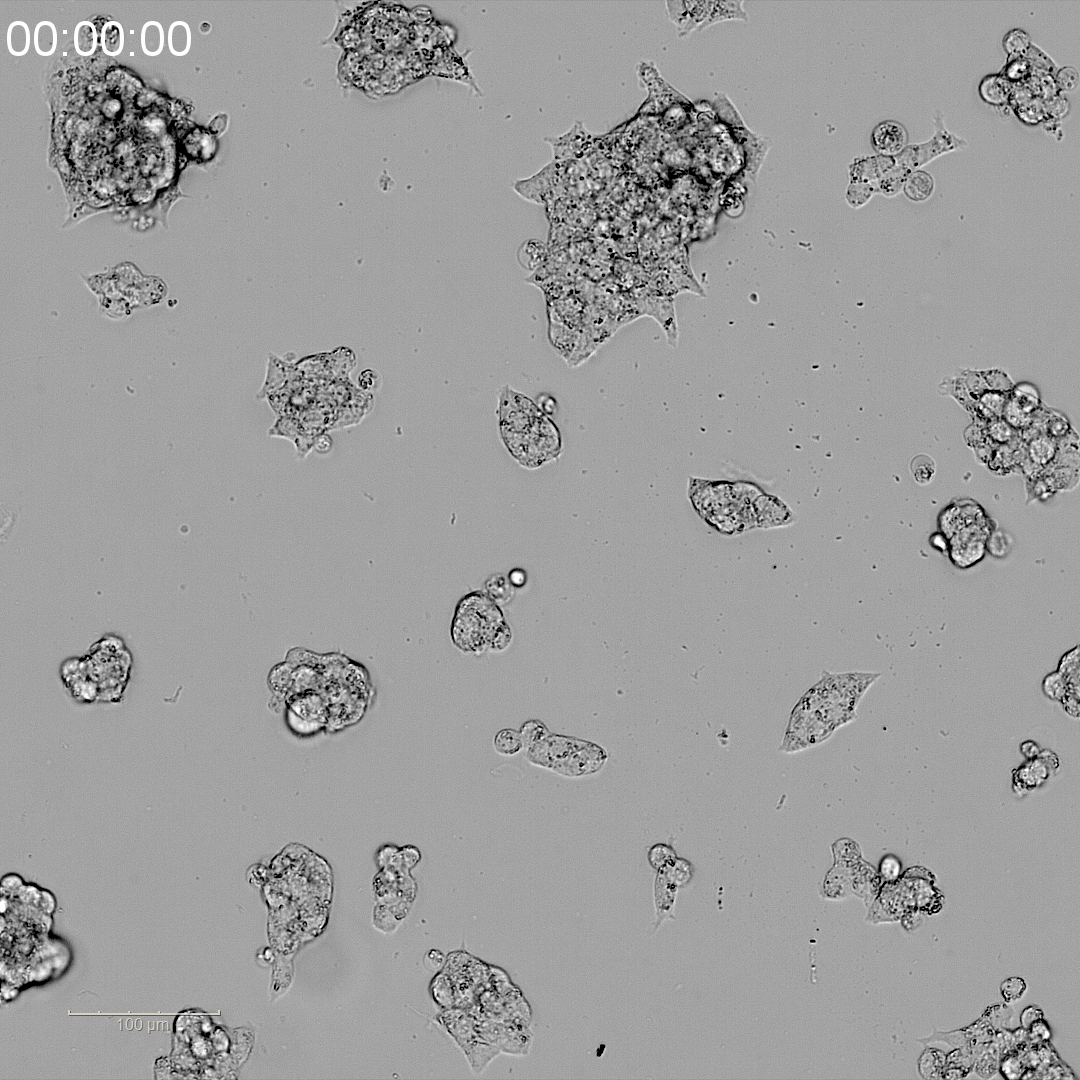

### Movie S2

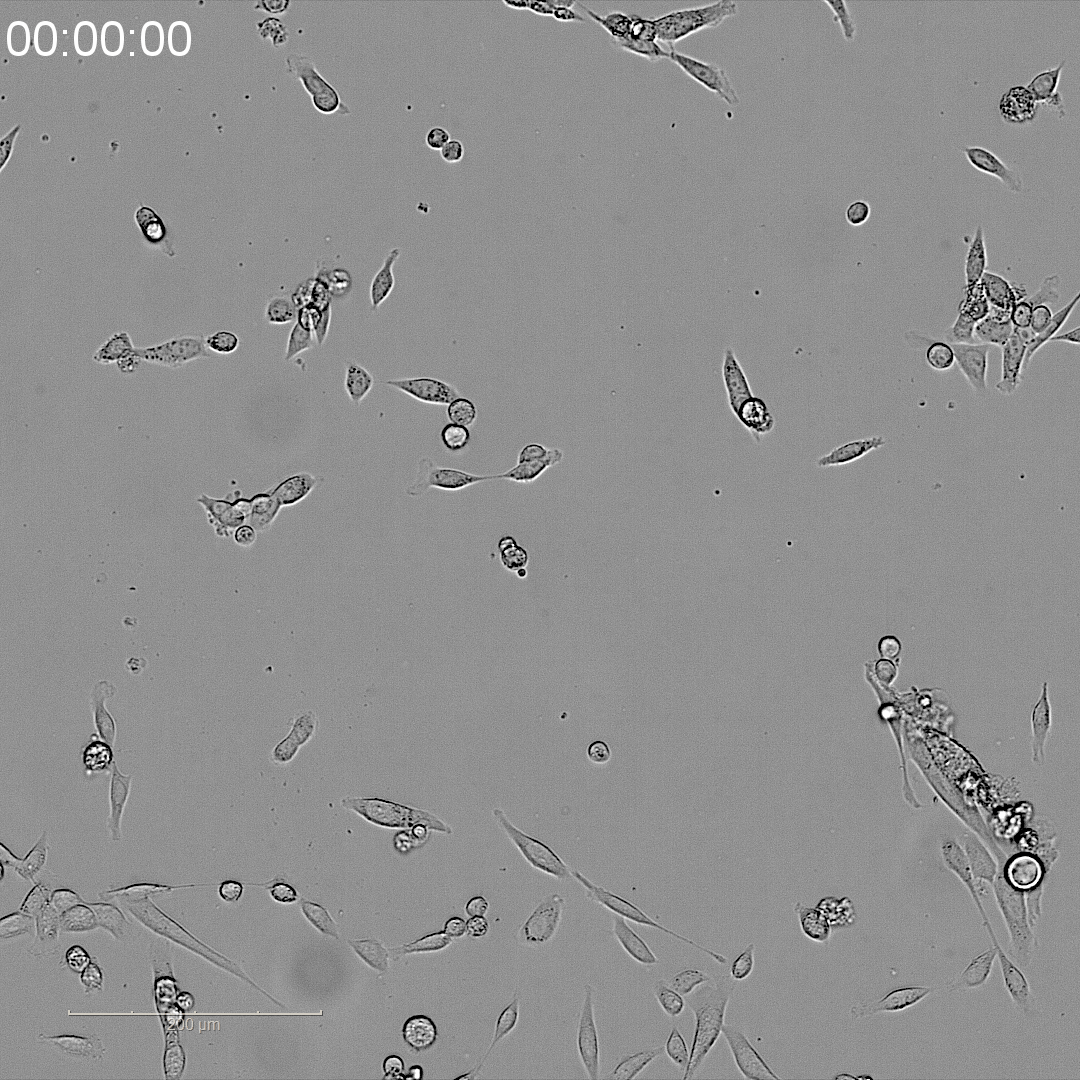

### Movie S3

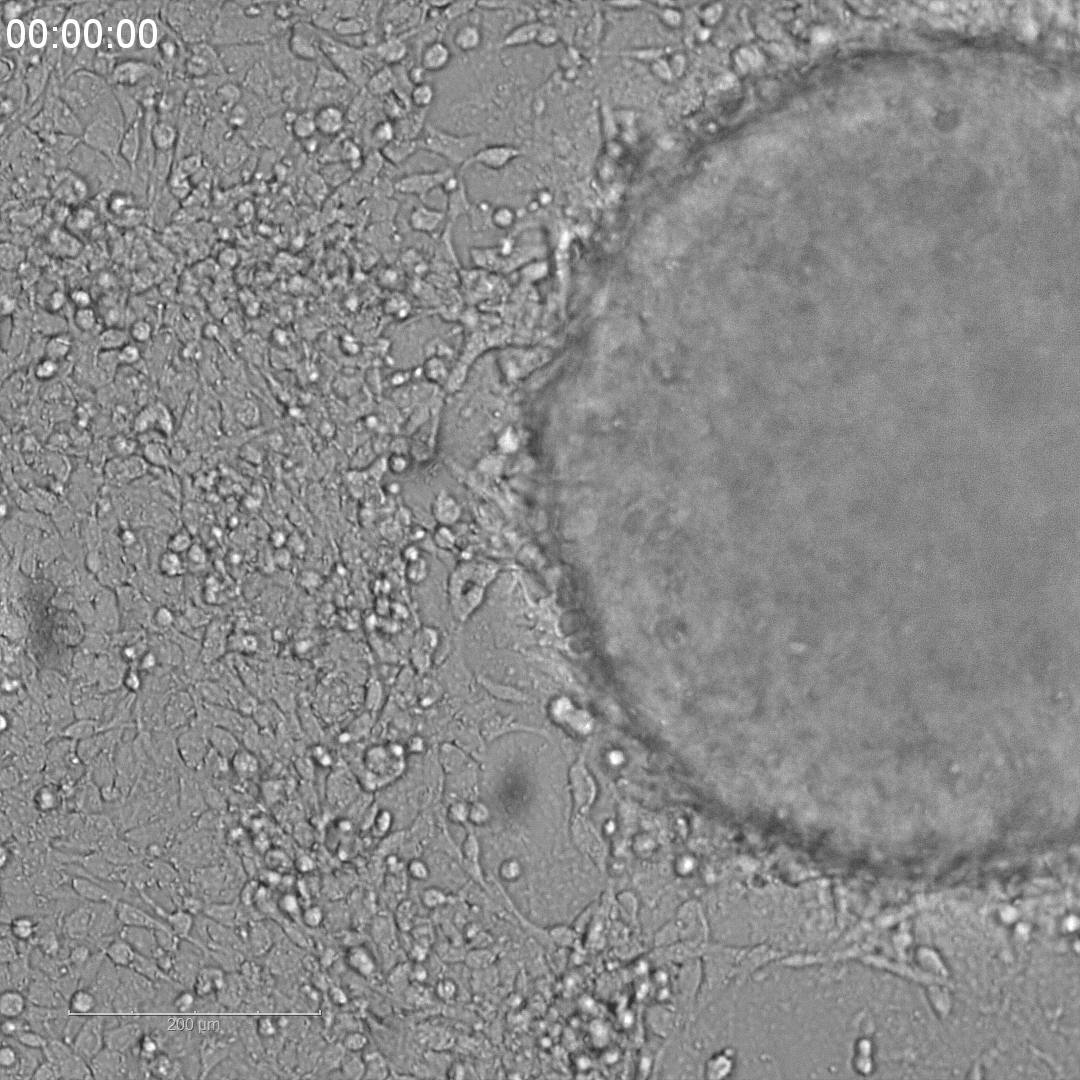
